# NeuroMechFly, a neuromechanical model of adult *Drosophila melanogaster*

**DOI:** 10.1101/2021.04.17.440214

**Authors:** Victor Lobato Ríos, Shravan Tata Ramalingasetty, Pembe Gizem Özdil, Jonathan Arreguit, Auke Jan Ijspeert, Pavan Ramdya

**Author notes:** equal contribution (Shravan Tata Ramalingasetty, Pembe Gizem Özdil).

## Abstract

Animal behavior emerges from a seamless interaction between neural network dynamics, musculoskeletal properties, and the physical environment. Accessing and understanding the interplay between these intertwined elements requires the development of integrative and morphologically realistic neuromechanical simulations. Until now, there has been no such simulation framework for the widely studied model organism, *Drosophila melanogaster*. Here we present NeuroMechFly, a data-driven model of the adult female fly within a physics-based simulation environment. NeuroMechFly combines a series of independent computational modules including a biomechanical exoskeleton with articulating body parts−legs, halteres, wings, abdominal segments, head, proboscis, and antennae−muscle models, and neural network controllers. To enable illustrative use cases, we first define minimal leg degrees-of-freedom by analyzing real 3D kinematic measurements during real *Drosophila* walking and grooming. Then, we show how, by replaying these behaviors using NeuroMechFly’s biomechanical exoskeleton in its physics-based simulation environment, one can predict otherwise unmeasured torques and contact reaction forces. Finally, we leverage NeuroMechFly’s full neuromechanical capacity to discover neural networks and muscle parameters that enable locomotor gaits optimized for speed and stability. Thus, NeuroMechFly represents a powerful testbed for building an understanding of how behaviors emerge from interactions between complex neuromechanical systems and their physical surroundings.

## 1 Introduction

Uncoupling the contributions to behavior of many neuronal and biomechanical elements is daunting. Systems-level numerical simulations can assist in this ambitious goal by consolidating data into a dynamic framework, generating predictions to be tested, and probing the sufficiency of prevailing theories to account for experimental observations [1–6]. Computational models, including neuromechanical simulations, have long played a particularly important role in the study of movement control in vertebrates [7–10] and invertebrates, including stick insects [11–14], cockroaches [15, 16], praying mantises [17], and ants [18].

For animals like invertebrates with a relatively small number of neurons that can be identified across individuals, a mapping of real to simulated biomechanical or circuit elements might enable a cross-talk whereby models make predictions that can then be tested experimentally. However, for many of the animals for which neuromechanical models currently exist, there is a dearth or absence of genetic tools that would facilitate repeatedly recording, or perturbing the same neurons across animals. By contrast, for a few commonly studied ‘model’ organisms, a dialogue between experimental results and computational predictions represents an exciting but largely unrealized opportunity. This is recently enabled by advances in computing power, the realism of physics-based simulation environments, and improvements in numerical optimization approaches. Neuromechanical models of some commonly studied organisms have already been developed including for the worm (*Caenorhabditis elegans* [19, 20]), maggots (larval *Drosophila melanogaster* [21]), and rodents [22]. However, for the adult fly, *Drosophila melanogaster*, only 2-dimensional (2D) [23] and morphologically unrefined [24] neuromechanical models exist.

Adult flies are an ideal organism for establishing a synergy between experimental and computational neuroscience. First, flies generate a large repertoire of complex behaviors including grooming [25], courtship [26], flight [27], and walking [28, 29] which they use to navigate complex environments [30]. The kinematics of these behaviors can now be quantified precisely using deep learning-based computer vision tools [31, 32] in 3-dimensions (3D) [33, 34]. Second, flies have a relatively small number of neurons that can be repeatedly genetically targeted [35] for recordings or perturbations in tethered, behaving animals [36–39]. These neurons can also be placed within their circuit context using recently acquired brain and ventral nerve cord (VNC) connectomes [40, 41]. We previously developed a simple physics-based simulation of adult *Drosophila melanogaster* to investigate hexapod locomotor gaits [24]. However, this older model has a number of important limitations that restrict its widespread use: it lacks (i) the morphological accuracy needed to simulate mass distributions, compliance, and physical constraints, (ii) muscle models and their associated passive dynamical properties, as well as (iii) neural networks or other control architectures.

Here we describe NeuroMechFly, a neuromechanical model of adult *Drosophila* that fills this methodological gap by incorporating a new, open-source computational framework consisting of exchangeable modules which provide access to biomechanics, neuromuscular control, and parameter optimization approaches. These modules maintain the capacity for whole organism simulation while also facilitating further open source extensions and improvements by the scientific community. Thus, NeuroMechFly is a completely new modeling framework and not simply an improvement of an earlier model [24].

The biomechanical exoskeleton of NeuroMechFly was obtained from a detailed CT-scan of an adult female fly which was then digitally rendered. We defined the model’s leg degrees-of-freedom based on an investigation of *Drosophila* 3D leg kinematics (Figure 1*A*), allowing us to discover that a previously unreported coxa-trochanter leg degree-of-freedom (DoF) is required to accurately recapitulate real fly walking and grooming. Using this biomechanical exoskeleton and replaying experimental leg kinematics within the PyBullet physics-based simulation environment (Figure 1*B*) [42], we then explored how one can estimate quantities that cannot be experimentally measured in behaving flies− ground reaction forces (GRFs), joint torques, and tactile contacts. As a second use-case illustration of NeuroMechFly’s potential, we leveraged the full neuromechanical framework−now including neural and muscle models−to show how the parameters of a central pattern generator (CPG)-inspired coupled-oscillator network and associated torsional spring and damper muscle model could be optimized to discover and explore controllers for fast and stable walking (Figure 1*C*). Importantly, the NeuroMechFly framework is modular and open-source, enabling future extensions including the use of more detailed neural and muscle models that permit more interpretable experimental predictions that can inform our understanding of real *Drosophila* neural circuits. Thus, NeuroMechFly represents an important step towards comprehending how behaviors emerge from a complex interplay between neural dynamics, musculoskeletal biomechanics, and physical interactions with the environment.

**Figure 1:**
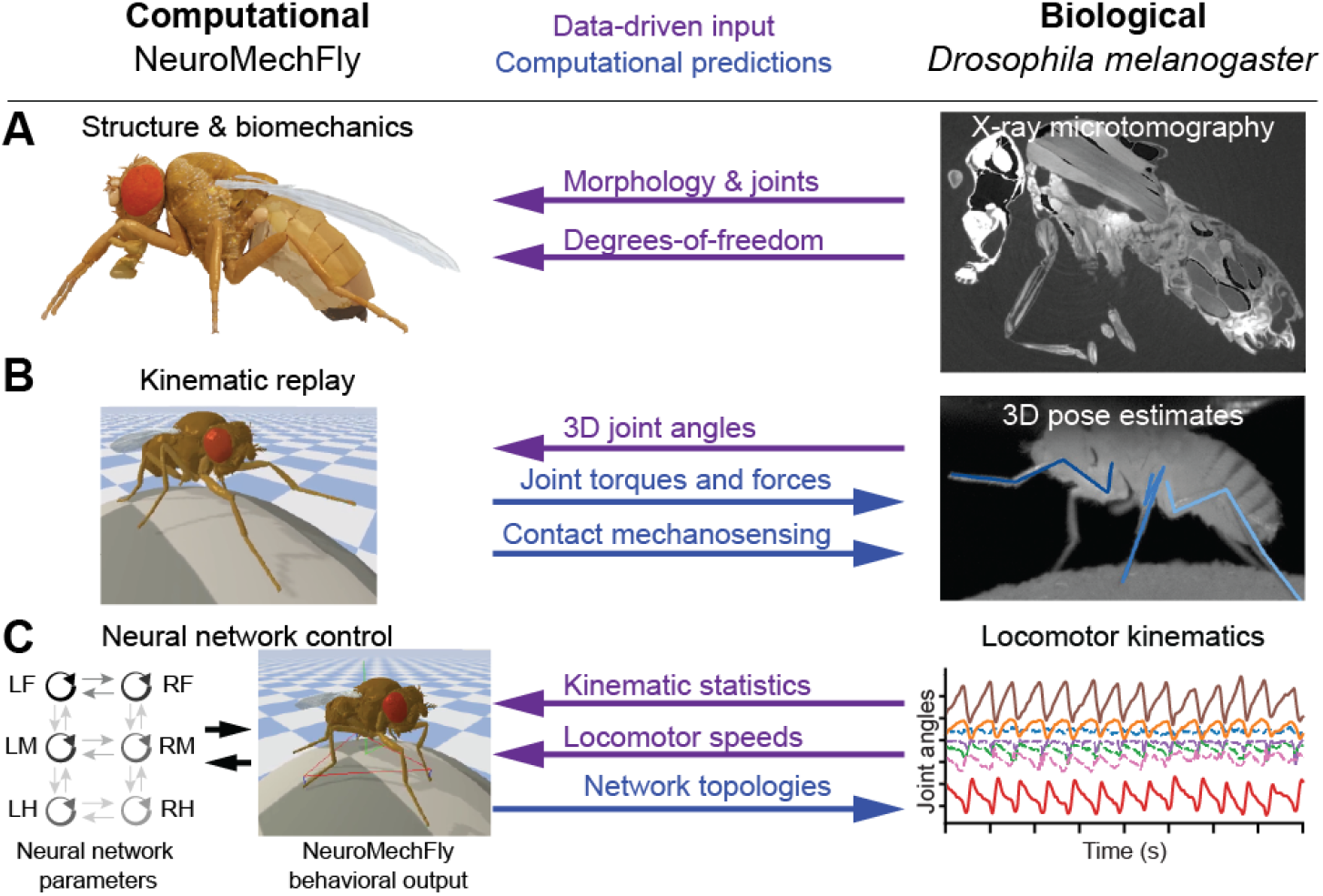
Data-driven development and applications of NeuroMechFly. **(A)** Body structures−morphology, joint locations, and degrees-of-freedom−were defined by x-ray microtomography and kinematic measurements. **(B)** Real 3D poses were used to replay kinematics in the model permitting the prediction of unmeasured contact reaction forces and joint torques. **(C)** Real limb kinematics were used to constrain the evolutionary optimization of neuromuscular parameters aiming to satisfy high-level objectives for walking−speed and static stability. The properties of optimized networks could then be more deeply analyzed.

## 2 Results

### 2.1 Constructing a data-driven biomechanical model of adult *Drosophila*

Behavior depends heavily on the body’s physical constraints and its interactions with the environment. Therefore, morphological realism is critical to accurately model leg movements and their associated self-collisions, joint ranges of motion, mass distributions, and mechanical loading. To achieve this level of realism in our model, we first measured the morphology of an adult female fly using x-ray microtomography **(Video 1)**. We first embedded the animal in resin to reduce blurring associated with scanner movements (Figure 2*A*). Then we processed the resulting microtomography data (Figure 2*B*) by binarizing it to discriminate between foreground (fly) and background (Figure 2*C*). Finally, we applied a Lewiner marching cubes algorithm [43] to generate a polygon mesh 3D reconstruction of the animal’s exoskeleton (Figure 2*D*).

**Figure 2:**
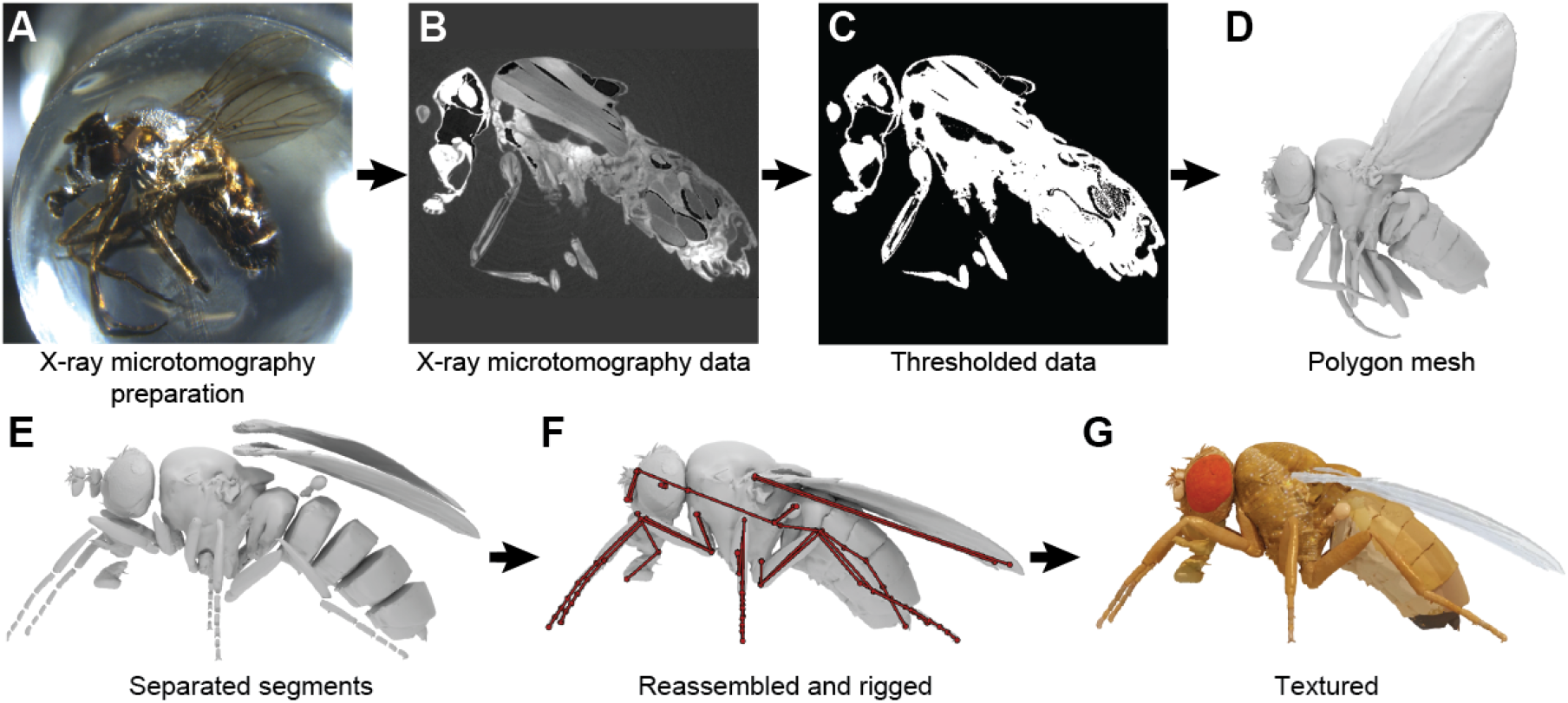
Constructing a data-driven biomechanical model of adult *Drosophila*. **(A)** An adult female fly is encased in resin for x-ray microtomography. **(B)** Cross-section of the resulting x-ray scan. Cuticle, muscles, nervous tissues, and internal organs are visible. **(C)** A threshold is applied to these data to separate the foreground (white) from the background (black). **(D)** A 3D polygon mesh of the exoskeleton and wings is constructed. **(E)** Articulated body parts are separated from one another. **(F)** These parts are reassembled into a natural resting pose. Joint locations are defined and constraints are introduced to create an articulated body (dark red). **(G)** Textures are added to improve the visual realism of the model.

Subsequently, to articulate appendages from this polygon mesh, we separated the body into 65 segments (see Table 1)(Figure 2*E*) and reassembled them into an empirically defined natural resting pose. Joints were added manually to permit actuation of the antennae, proboscis, head, wings, halteres, abdominal segments, and leg segments. Leg articulation points were based on observations from high-resolution videography [33], and previously reported leg DoFs [44–46](Table 1)(Figure 2*F*). By measuring leg segment lengths across animals (n = 10), we confirmed that the model’s legs are within the range of natural size variation (Figure S1).

**Table 1:** Model body parts and degrees-of-freedom between each segment and its parent.

| Body part | Segment | Parent | Degrees of freedom |
| --- | --- | --- | --- |
| Abdomen | A1A2 | Thorax | 1 |
|  | A3 | A1A2 | 1 |
|  | A4 | A3 | 1 |
|  | A5 | A4 | 1 |
|  | A6 | A5 | 1 |
| Head | Head capsule | Thorax | 3 |
|  | Eyes (x2) | Head | 0 |
|  | Antennae (x2) |  | 1 |
|  | Rostrum |  | 1 |
|  | Haustellum | Rostrum | 1 |
| Legs | Coxa (x6) | Thorax | 3 |
|  | Trochanter/Femur (x6) | Coxa | 2 |
|  | Tibia (x6) | Femur | 1 |
|  | Tarsus1 (x6) | Tibia | 1 |
|  | Tarsus2 (x6) | Tarsus1 | 1 |
|  | Tarsus3 (x6) | Tarsus2 | 1 |
|  | Tarsus4 (x6) | Tarsus3 | 1 |
|  | Tarsus5-Claw (x6) | Tarsus4 | 1 |
| Thorax | Halteres (x2) | Thorax | 3 |
|  | Wings (x2) |  | 3 |
|  | Thorax | - | 0 |

To facilitate the control of each DoF in the physics engine, we used hinge-type joints to connect each of the body parts. We later show that this approximation permits accurate replay of leg end-effector trajectories. Therefore, to construct thorax-coxa joints with three DoFs, we combined three hinge joints along the yaw, pitch, and roll axes of the base link. Finally, we textured the model for visualization purposes (Figure 2*G*). This entire process yielded a rigged model of adult *Drosophila* with the morphological accuracy required for biomechanical studies as well as, in potential future work, model-based computer vision tasks like pose estimation [47–51].

### 2.2 Identifying minimal joint degrees-of-freedom required to accurately replay real 3D leg kinematics

After constructing an articulating biomechanical model of an adult fly, we next asked whether the six reported and implemented leg DoFs−(i-iii) thorax-coxa (ThC) elevation/depression, protraction/retraction, and rotation, (iv) coxa-trochanter (CTr) flexion/extension, (v) femur-tibia (FTi) flexion/extension, and (vi) tibia-tarsus (TiTa) flexion/extension [44, 45]−would be sufficient to accurately replay measured 3D leg kinematics. We did not add a trochanter-femur (TrF) joint because the *Drosophila* trochanter is thought to be fused to the femur [45]. For the middle and hind legs, ThC protraction/retraction occurs along a different axis than similarly named movements of the front legs.

Therefore, we chose to instead use the notations ‘roll’, ‘pitch’, and ‘yaw’ to refer to rotations around the anterior/posterior, medial/lateral, and dorsal/ventral axes of articulated segments, respectively (**Video 2**).

For our studies of leg kinematics, we focused on forward walking and grooming, two of the most common spontaneously-generated *Drosophila* behaviors. First, we used DeepFly3D [33] to acquire 3D poses from recordings of tethered flies behaving spontaneously on a spherical treadmill. Due to 3D pose estimation-related noise and some degree of inter-animal morphological variability (Figure S1), directly actuating NeuroMechFly using raw 3D poses was impossible. To overcome this issue, we fixed the positions of base ThC joints as stable reference points and set each body part’s length to its mean length for a given experiment. Then, we scaled relative ThC positions and body part lengths using our biomechanical model as a template. Thus, instead of using 3D cartesian coordinates, we could now calculate joint angles that were invariant across animals and that matched the DoFs used by NeuroMechFly. At first we calculated these joint angles for the six reported DoFs [44, 45] by computing the dot product between the global rotational axes and coxal joints and between adjacent leg segments joined by single-rotational joints (see Materials and Methods).

When only these six DoFs were used to replay walking and grooming, we consistently observed a large discrepancy between 3D pose-derived cartesian joint locations and those computed from joint angles via forward kinematics (Figure 3, Base DoF Dot product). Visualization of these errors showed significant out-of-plane movements of the tibia and tarsus (**Video 3**, top-left). This was surprising given that each leg is thought to consist of a ball-and-socket joint (three DoFs in the ThC joint) followed by a series of one DoF hinge joints that, based on their orientations, should result in leg segments distal to the coxa residing in the same plane. Therefore, we next tried to identify alternative leg configurations that might better match 3D poses. First we performed an inverse kinematics optimization of joint angles rather than dot product operations. This would allow us to identify angle configurations that minimize error at the most distal tip of the kinematic chain−in this case, the pretarsus. Although inverse kinematics yielded a lower discrepancy (Figure 3, Base DoF Inverse kinematics), we still observed consistent out-of-plane leg movements (**Video 3**, top-middle).

**Figure 3:**
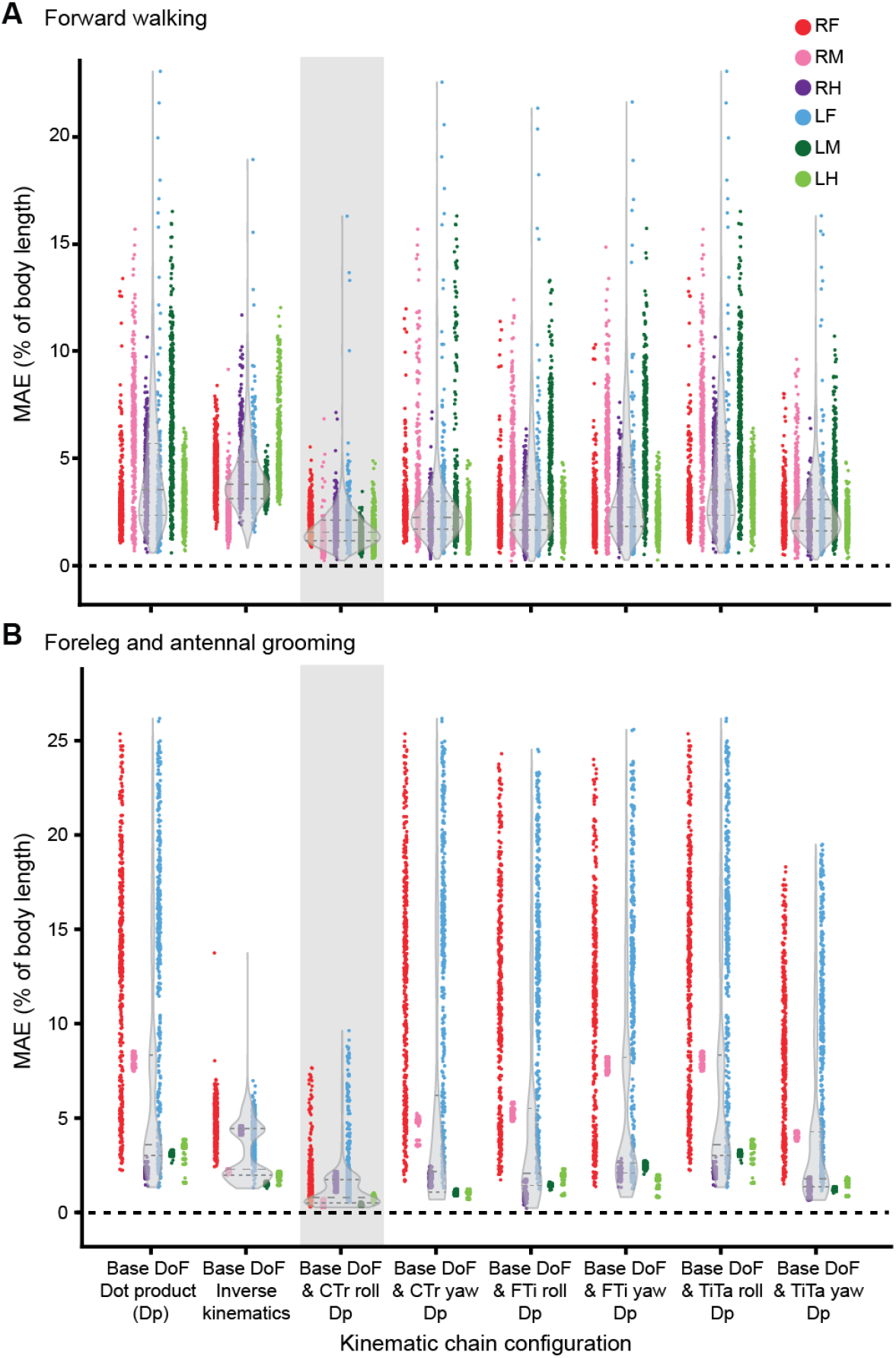
Adding a CTr roll DoF to base DoFs enables the most accurate kinematic replay of real walking and grooming. Body-length normalized mean absolute errors (MAE) comparing measured 3D poses and angle-derived joint positions for various DoF configurations. Measurements were made for representative examples of **(A)** forward walking, or **(B)** foreleg/antennal grooming. For each condition, n = 2400 samples were computed for all six legs across 4 s of 100 Hz video data. Data for each leg are color-coded. ‘R’ and ‘L’ indicate right and left legs, respectively. ‘F’, ‘M’, and ‘H’ indicate front, middle, and hind legs, respectively. Violin plots indicate median, upper, and lower quartiles (dashed lines). Results from adding a coxa-trochanter roll DoF to based DoFs are highlighted in light gray.

We next examined whether an extra DoF might be needed at the CTr joint to accurately replicate real fly leg movements. This analysis was motivated by the fact that: (i) other insects use additional stabilizing rotations at or near the TrF joint [52–55], (ii) unlike other insects, the *Drosophila* trochanter and femur are fused, and (iii) *Drosophila* hosts reductor muscles of unknown function near the CTr joint [44]. To ensure that any improvements did not result simply from overfitting by increasing the number of DoFs, we also tested the effect of adding one roll or yaw DoF to each of the more distal hinge-type joints (CTr, FTi and TiTa)(**Video 2**). Indeed, for both walking (**Video 3**, top-right) and foreleg/antennal grooming (**Video 4**, top-right), we observed that adding a CTr roll DoF to the six previously reported (‘base’) DoFs significantly and uniquely reduced the discrepancy between 3D pose-derived and forward kinematics-derived joint positions, even when compared with improvements from inverse kinematics (Figure 3, Base DoF & CTr roll; for statistical analysis, see Table 2 and Table 3). This improvement was also evident on a joint-by-joint basis for walking (Figure S2) and grooming (Figure S3) and it was not achieved by any other kinematic chain tested−a result that argues against the possibility of over-fitting (Figure 3, Base DoF & CTr yaw, Base DoF & FTi roll, Base DoF & FTi yaw, Base DoF & TiTa roll, Base DoF & TiTa yaw). These findings demonstrate that accurate kinematic replay of *Drosophila* leg movements requires seven DoFs per leg: the previously reported six DoFs [44, 45] as well as a roll DoF near the CTr joint. Thus, by default, NeuroMechFly’s biomechanical exoskeleton incorporates this additional DoF for each leg (Table 1).

**Table 2:** Matrix of p-values from pairwise comparisons of position errors after calculating forward kinematics for walking. Numbers in bold (except in the case of identity) indicate that the p-value *>* 0.001 (i.e., no statistical difference).

|  | Base | IK | Base & CTr roll | Base & CTr yaw | Base & FTi roll | Base & FTi yaw | Base & TiTa roll | Base & TiTa yaw |
| --- | --- | --- | --- | --- | --- | --- | --- | --- |
| Base | 1.00 | 5.42e-13 | 0.00 | 7.08e-184 | 2.28e-133 | 4.53e-50 | **9.95e-01** | 1.53e-197 |
| IK | 5.42e-13 | 1.00 | 0.00 | 4.48e-285 | 4.37e-222 | 6.82e-110 | 5.42e-13 | 8.62e-302 |
| Base & CTr roll | 0.00 | 0.00 | 1.00 | 5.49e-138 | 2.96e-189 | 0.00 | 0.00 | 1.57e-126 |
| Base & CTr yaw | 7.08e-184 | 4.48e-285 | 5.49e-138 | 1.00 | 2.52e-05 | 5.13e-45 | 7.83e-184 | **5.38e-01** |
| Base & FTi roll | 2.28e-133 | 4.37e-222 | 2.96e-189 | 2.52e-05 | 1.00 | 8.33e-22 | 2.44e-133 | 1.08e-07 |
| Base & FTi yaw | 4.53e-50 | 6.82e-110 | 0.00 | 5.13e-45 | 8.33e-22 | 1.00 | 4.53e-50 | 6.05e-52 |
| Base & TiTa roll | **9.95e-01** | 5.42e-13 | 0.0 | 7.83e-184 | 2.44e-133 | 4.53e-50 | 1.00 | 1.71e-197 |
| Base & TiTa yaw | 1.53e-197 | 8.63e-302 | 1.57e-126 | **5.38e-01** | 1.08e-07 | 6.05e-52 | 1.71e-197 | 1.00 |

**Table 3:** Matrix of p-values matrix from pairwise comparisons of position errors after calculating forward kinematics for grooming. Numbers in bold (except in the case of identity) indicate that the p-value *>* 0.001 (i.e., no statistical difference).

|  | Base | IK | Base & CTr roll | Base & CTr yaw | Base & FTi roll | Base & FTi yaw | Base & TiTa roll | Base & TiTa yaw |
| --- | --- | --- | --- | --- | --- | --- | --- | --- |
| Base | 1.00 | 4.34e-128 | 0.00 | 7.57e-149 | 2.59e-131 | 4.72e-32 | **1.00** | 2.47e-192 |
| IK | 4.34e-128 | 1.00 | 0.00 | **2.02e-01** | **1.00** | 4.30e-34 | 3.27e-126 | 1.11e-07 |
| Base & CTr roll | 0.00 | 0.00 | 1.00 | 0.00 | 0.00 | 0.00 | 0.00 | 0.00 |
| Base & CTr yaw | 7.57e-149 | **2.02e-01** | 0.00 | 1.00 | **3.04e-01** | 2.56e-45 | 8.05e-147 | **1.08e-03** |
| Base & FTi roll | 2.59e-131 | **1.00** | 0.00 | **3.04e-01** | 1.00 | 8.96e-36 | 2.08e-129 | 5.70e-07 |
| Base & FTi yaw | 4.72e-32 | 4.30e-34 | 0.00 | 2.56e-45 | 8.96e-36 | 1.00 | 3.84e-31 | 4.86e-71 |
| Base & TiTa roll | **1.00** | 3.27e-126 | 0.00 | 8.05e-147 | 2.08e-129 | 3.84e-31 | 1.00 | 4.85e-190 |
| Base & TiTa yaw | 2.47e-192 | 1.11e-07 | 0.00 | **1.08e-03** | 5.70e-07 | 4.86e-71 | 4.85e-190 | 1.00 |

### 2.3 Using NeuroMechFly to estimate joint torques and contact forces through kinematic replay of real fly behaviors

Having identified a suitable set of leg DoFs, we next aimed to illustrate the utility of NeuroMechFly as a biomechanical model within the PyBullet physics-based environment. PyBullet is an integrative framework that not only gives access to collisions, reaction forces, and torques but also imposes gravity, time, friction, and other morphological collision constraints, allowing one to explore their respective roles in observed animal behaviors. Specifically, we focused on testing the extent to which one might use kinematic replay of real behaviors to infer torques, and contact forces like body part collisions and ground reaction forces (GRFs)−quantities that remain technically challenging to measure in small insects like *Drosophila* [18,56]. Although kinematic replay may not provide information about internal forces that are not reflected in 3D poses (e.g., how tightly the legs grip the spherical treadmill without changes in posture), estimates of collisions and interaction forces may be a good first approximation of an animal’s proprioception and mechanosensation.

We explored this possibility by using a proportional-derivative (PD) controller implemented in PyBullet to actuate the model’s leg joints, replaying measured leg kinematics during forward walking and foreleg/antennal grooming. We used joint angles and angular velocities as target signals for the controller. Because, when applying this kind of controller, there is no unique set of contact solutions that match forces and torques to prescribed kinematics (i.e., experimental validation of force estimates would ultimately be necessary), we first quantified how sensitive torque and force estimates were to changes in PD controller gains. Based on this sensitivity analysis, we selected gain values that optimized the precision of kinematic replay (Figure S4, blue squares) and for which small deviations did not result in large variations in measured physical quantities (Figure S5, red traces). We included all seven leg degrees-of-freedom from our error analysis (Figure S6) and the model’s ‘zero-angle pose’ was selected to make joint angles intuitive (Figure S7). We also set fixed values for the orientation of abdominal segments, wings, halteres, head, proboscis, and antennae to generate a natural pose (Table 4).

**Table 4:** Fixed angles for body joints during kinematic replay and optimization.

| Body part | Joint | Fixed angle (deg) | Body part | Joint | Fixed angle (deg) |
| --- | --- | --- | --- | --- | --- |
| Abdomen | A1A2 | 0 | Thorax | Left haltere roll | 0 |
|  | A3 | -15 |  | Left haltere pitch | 0 |
|  | A4 | -15 |  | Left haltere yaw | 0 |
|  | A5 | -15 |  | Right haltere roll | 0 |
|  | A6 | -15 |  | Right haltere pitch | 0 |
| Head | Head capsule roll | 0 |  | Right haltere yaw | 0 |
|  | Head capsule pitch | 10 |  | Left wing roll | 90 |
|  | Head capsule yaw | 0 |  | Left wing pitch | 0 |
|  | Left antenna | 35 |  | Left wing yaw | -17 |
|  | Right antenna | -35 |  | Right wing roll | -90 |
|  | Rostrum | 90 |  | Right wing pitch | 0 |
|  | Haustellum | -60 |  | Right wing yaw | 17 |

**Table 5:** Fixed angles for leg joints during optimization (deg).

| Body Part | Side | ThC yaw | ThC pitch | ThC roll | CTr pitch | CTr roll | FTi | TiTa |
| --- | --- | --- | --- | --- | --- | --- | --- | --- |
| Front | Left | 0 | actuated | 10 | actuated | 0 | actuated | -39 |
|  | Right | 0 | actuated | -10 | actuated | 0 | actuated | -39 |
| Middle | Left | 7.45 | -5 | actuated | actuated | 0 | actuated | -54 |
|  | Right | -7.45 | -5 | actuated | actuated | 0 | actuated | -54 |
| Hind | Left | 3.45 | 6.2 | actuated | actuated | 0 | actuated | -45 |
|  | Right | -3.45 | 6.2 | actuated | actuated | 0 | actuated | -45 |

When we replayed walking (Figure 4*A-C*)(**Video 5**) and foreleg/antennal grooming (Figure 5*A-C*) (**Video 6**), we observed that the model’s leg movements were largely identical to those measured from *Drosophila*. By measuring real ball rotations [57] and comparing them with simulated spherical treadmill rotations, for a range of soft constraint parameters (Figure S8), we quantified high similarity between real and simulated spherical treadmill forward velocities (Figure S9*D*), and to some extent, yaw velocities (Figure S9*F*). Sideways velocities were smaller and, thus, difficult to compare (Figure S9*E*). This was notable given that the ball’s rotations were not explicitly controlled but emerged from tarsal contacts and forces in our simulation. These observations support the accuracy of our computational pipeline in processing and replaying recorded joint positions.

**Figure 4:**
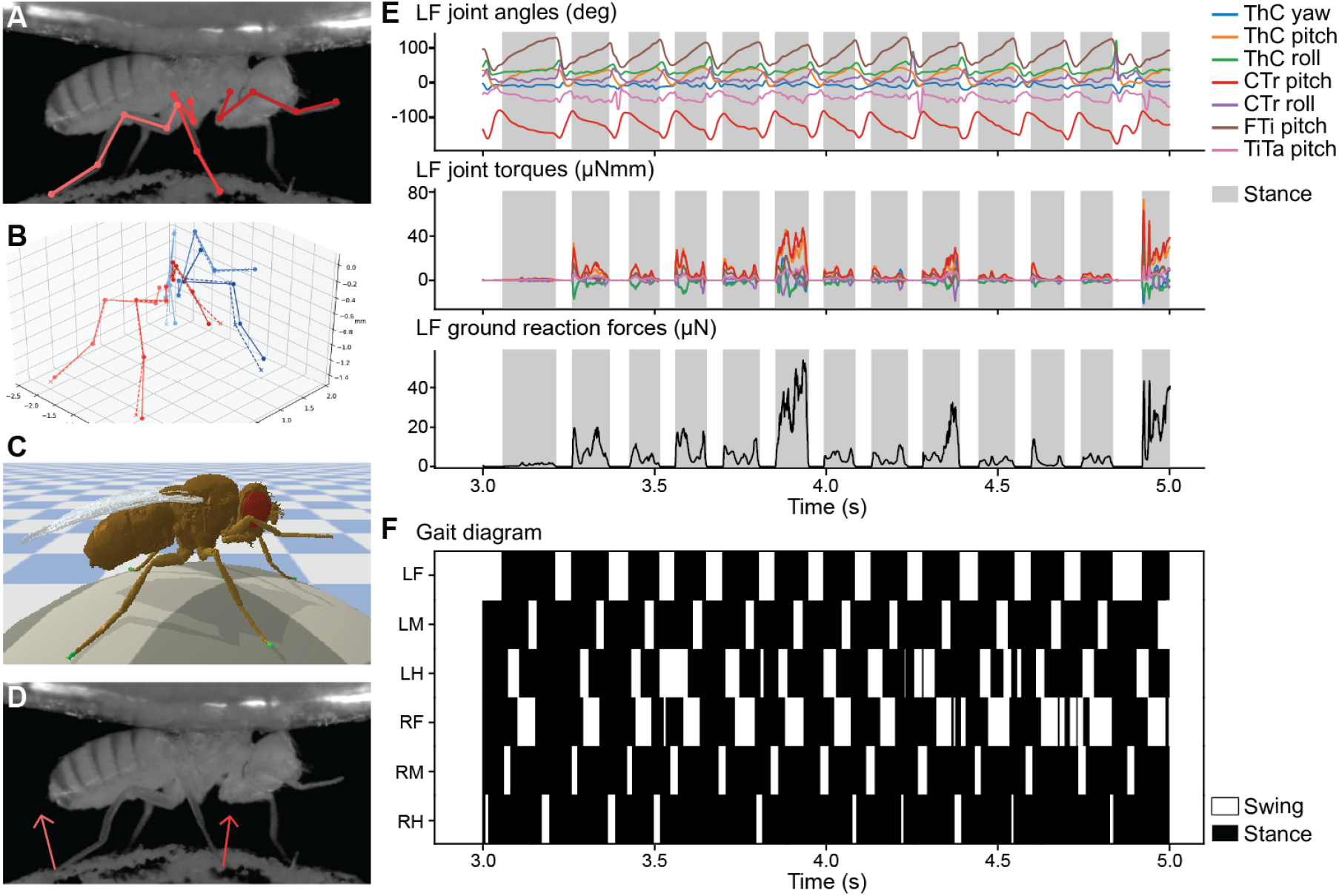
Kinematic replay of forward walking allows the estimation of ground contacts and reaction forces. **(A)** Multiple cameras and deep learning-based 2D pose estimation are used to track the positions of each leg joint while a tethered fly is walking on a spherical treadmill. **(B)** Multiview 2D poses (solid lines) are triangulated and processed to obtain 3D joint positions (dashed lines). These are further processed to compute joint angles for seven DoFs per leg. **(C)** Joint angles are replayed using PD control in NeuroMechFly. Body segments in contact with the ground are indicated (green). **(D)** Estimated ground reaction force vectors (red arrows) are superimposed on original video data. **(E, top)** Kinematic replay of real 3D joint angles permits estimation of unmeasured **(E, middle)** joint torques, and **(E, bottom)** ground reaction forces. Only data for the left front leg (LF) are shown. Grey bars indicate stance phases when the leg is in contact with the ground. Joint DoFs are color-coded. **(F)** A gait diagram illustrating stance (black) and swing (white) phases for each leg as computed by measuring simulated tarsal contacts with the ground.

**Figure 5:**
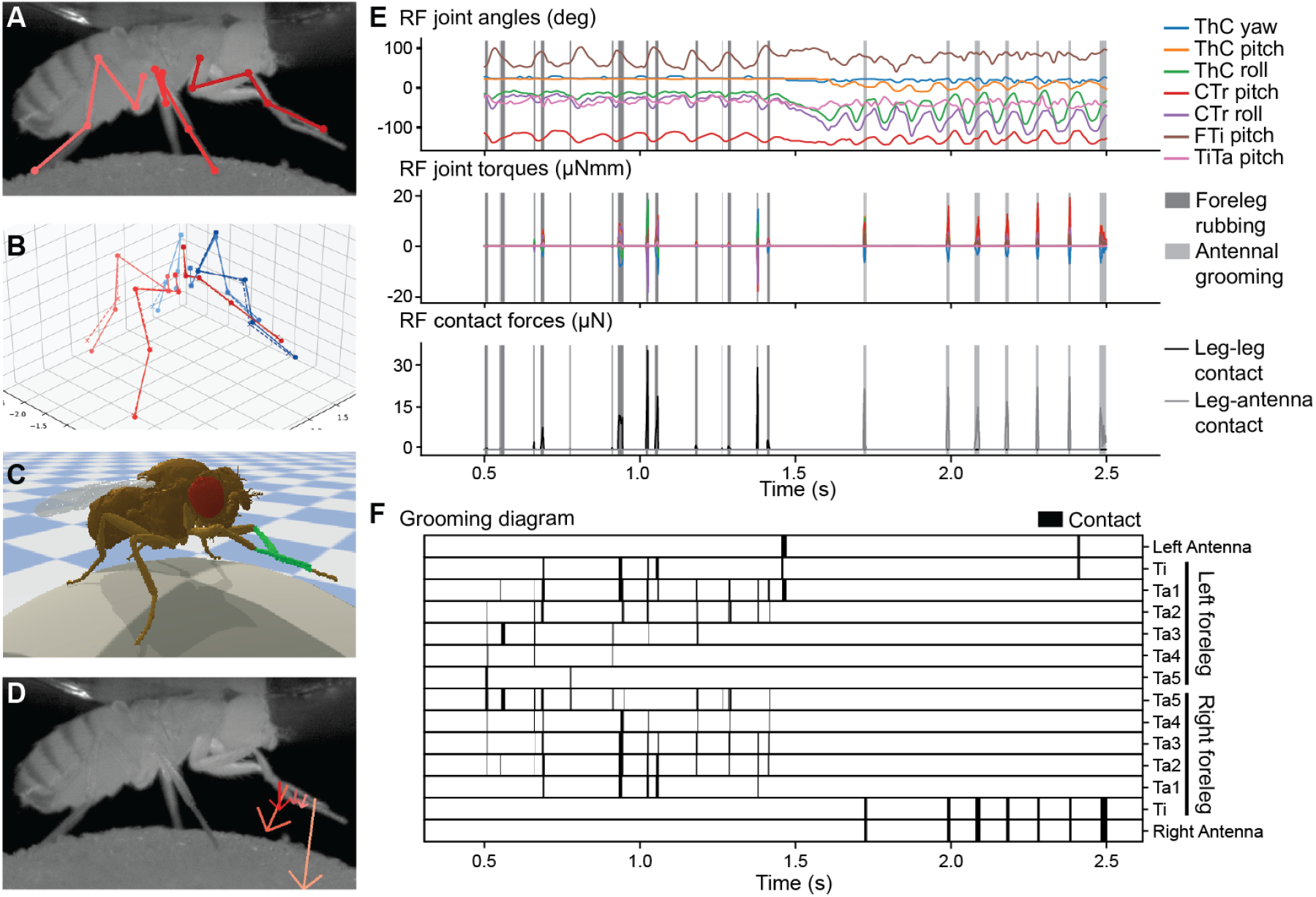
Kinematic replay allows the estimation of self-collisions and reaction forces during foreleg/antennal grooming. **(A)** Multiple cameras and deep learning-based 2D pose estimation are used to track the positions of each leg joint while a tethered fly grooms its forelegs and antennae. **(B)** Multiview 2D poses (solid lines) are triangulated and processed to obtain 3D joint positions (dashed lines). These are further processed to compute joint angles for seven DoFs per leg. **(C)** Joint angles are replayed using PD control in NeuroMechFly. Body segments undergoing collisions are indicated (green). **(D)** Estimated leg-leg and leg-antennae contact forces (red arrows) are superimposed on original video data. **(E, top)** Kinematic replay of real joint angles permits estimations of unmeasured **(E, middle)** joint torques, and **(E, bottom)** contact forces. Only data for the right front (RF) leg are shown. Dark grey bars indicate leg-leg contacts. Light grey bars indicate leg-antenna contacts. Joints are color-coded. **(F)** A grooming diagram illustrating contacts (black) made by the front leg’s five tarsal segments (‘Ta1’ and ‘Ta5’ being the most proximal and the most distal, respectively), tibia (‘Ti’), and both antennae (‘Ant’).

Next, we more directly validated collisions and forces computed within the PyBullet physics-based simulation environment. From kinematic replay of joint angles during walking (Figure 4*E*, top), we measured rich, periodic torque dynamics (Figure 4*E*, middle). These were accompanied by ground reaction forces (GRFs) that closely tracked subtle differences in leg placement across walking cycles (Figure 4*E*, bottom). Superimposing these GRF vectors on raw video recordings of the fly allowed us to visualize expected tarsal forces (Figure 4*D*)(**Video 5**, top-left) which could also be used to generate predicted gait diagrams during tethered walking (Figure 4*F*). These predictions were highly accurate (83.5 - 87.3% overlap) when compared with manually labeled ground-truth gait diagrams for three different animals and experiments (Figure S10). This result was notable given that the thorax is fixed and, in principle, subtle changes in attachment height could increase or decrease the duration of leg-treadmill contacts.

Similarly, for foreleg/antennal grooming (Figure 5*A-C*), we observed that measured joint angles (Figure 5*E*, top) could give rise to complex torque dynamics (Figure 5*E*, middle). Associated leg and antennal contact forces (Figure 5*D, E*, bottom) reached magnitudes about three times the fly’s weight. These fall within the range of previously observed maximum forces measured at the tip of the tibia (~ 100*µN*) for ballistic movements [58], but further experimental data will be required to fully validate these measurements. These leg and antennal contact forces were used to generate groom-ing diagrams−akin to locomotor gait diagrams−that illustrate predicted contacts between distal leg segments and the antennae (Figure 5*F*). During leg-leg grooming, we observed collisions that moved continuously along the leg segments in proximal to distal sweeps. These collision data provide a richer description of grooming beyond classifying the body part that is being cleaned and can enable a more precise physical quantification of many other behaviors including, for example, inter-animal boxing or courtship tapping. This approach also revealed the importance of having a morphologically accurate biomechanical model. When we replaced our CT scan-based leg segments and antennae with more conventional stick segments having similar diameters and lengths, we observed less rich collision dynamics including the elimination of interactions between the tarsi and antennae (Figure S11) **(Video 7)**.

Because our 3D pose estimates were made on a tethered fly behaving on a spherical treadmill, we also ‘tethered’ our simulation by fixing the thorax position. Next, we asked to what extent our model might be able to walk without body support (i.e., keeping its balance while carrying its body weight). To do this, we replayed 3D kinematics from tethered walking (Figure 4)(**Video 5**) while NeuroMechFly could walk freely (untethered) on flat terrain. Indeed, we observed that our model walked stably on the ground (**Video 8**). Although an animal’s legs would naturally be positioned differently on a curved versus a flat surface, the flexibility of NeuroMechFly’s tarsal segments allowed it to walk freely with a natural pose using 3D poses taken from tethered walking on a curved spherical treadmill. As expected, flat ground locomotion matched the velocities of tethered walking (Figure S12) better than walking paths (**Video 8**): small deviations in heading direction yield large changes in trajectories.

In summary, we have shown how NeuroMechFly’s biomechanical exoskeleton−without muscle or neuron models−can be used to replay real 3D poses to estimate otherwise inaccessible physical quantities like joint torques, collisions, and reaction forces that are accessible from its physics-based simulation engine.

### 2.4 Using NeuroMechFly to explore locomotor controllers by optimizing CPG-oscillator networks and muscles

As a full neuromechanical model, NeuroMechFly consists not only of biomechanical elements, like those used for kinematic replay, but also neuromuscular elements. In our computational framework, these represent additional modules that the investigator can define to be more abstract−e.g., leaky integrate-and-fire neurons and spring-and-damper models−or more detailed−e.g., Hodgkin-Huxley neurons and Hill-type muscle models. Parameters for neural networks and muscles that maximize user-defined objectives and minimize penalties can be identified using evolutionary optimization.

Here, to provide a proof-of-concept of this approach, we aimed to discover neuromuscular controllers that optimize fast and statically stable tethered walking. Insect walking gaits are commonly thought to emerge from the connectivity and dynamics of networks of CPGs within the ventral nerve cord (VNC) [15, 16, 59, 60]. Although alternative, decentralized approaches have also been proposed [14, 61], we focused on exploring a CPG-based model of locomotor control. First, we designed a neural network controller consisting of a CPG-like coupled oscillator [62] for each joint (Figure 6*A*). For simplicity, we denote the output of each coupled oscillator as the activity of a CPG. These CPGs, in turn, were connected to spring-and-damper (‘Ekeberg-type’) muscles [63]. This simple muscle model has been used to effectively simulate lamprey [63], stick insect [11], and salamander [9] locomotion.

**Figure 6:**
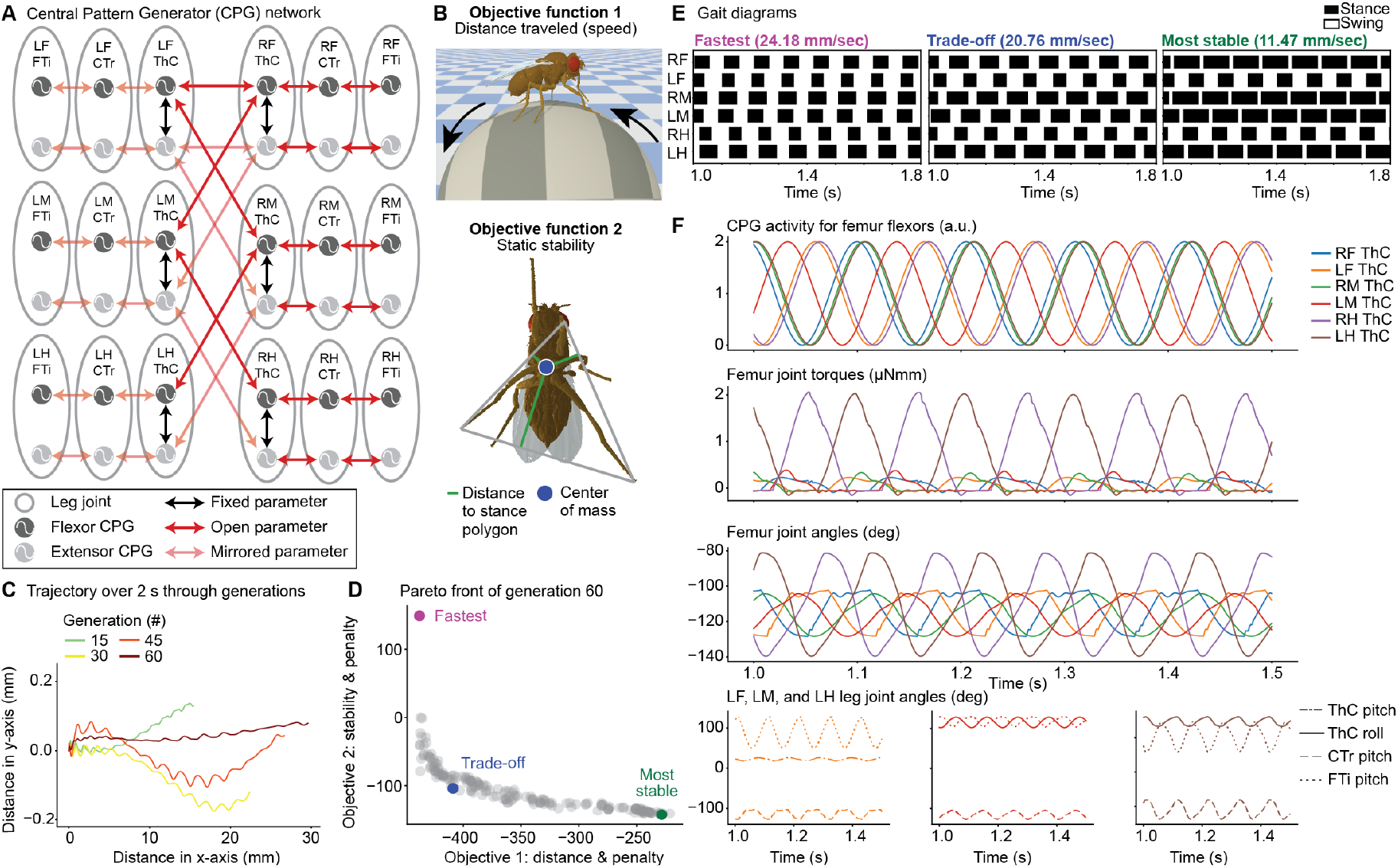
Using evolutionary optimization to identify oscillator network and muscle parameters that achieve fast and stable locomotion. **(A)** A network of coupled oscillators modeling CPG-based intra- and interleg circuits in the ventral nerve cord of *Drosophila*. Oscillator pairs control specific antagonistic leg DoFs (gray). Network parameter values are either fixed (black), modified during optimization (red), or mirrored from oscillators on the other side of the body (pink). **(B)** Multi-objective optimization of network and muscle parameters maximizes forward walking distance traveled (speed) and static stability. **(C)** A ‘trade-off’ solution’s locomotor trajectory (distance traveled over x and y axes) across 60 optimization generations. **(D)** Pareto front of solutions from the final (60th) optimization generation. Three individuals were selected from the population using different criteria: the longest distance traveled (fastest, purple), the most statically stable solution (‘most stable’, green), and the solution having the smallest 2-norm of both objective functions after normalization (trade-off). **(E)** Gait diagrams for selected solutions from generation 60. Stance (black) and swing (white) phases were determined based on tarsal ground contacts for each leg. Velocity values were obtained by averaging the ball’s forward velocity over 2 s. **(F)** Central Pattern Generator (CPG) outputs, joint torques, and joint angles of each leg’s femur for the ‘trade-off’ solution. Intraleg joint angles for the left front, middle, and hind legs are also shown. Legs are color-coded and joints are shown in different line styles.

We aimed to identify suitable neuromuscular parameters for walking in an reasonably short period of optimization time (less than 24 h per run on a workstation). Therefore, we reduced the number of parameters and, thus, the search space. Specifically, we limited controlled DoFs to those which (i) were sufficient to generate walking in other insect simulations [64] and (ii) had the most pronounced effect on overall leg trajectories in our kinematic analysis of real flies (Figure S13). Thus, we used the following three DoFs per leg that satisfied these criteria: CTr pitch, and FTi pitch for all legs as well as ThC pitch for the forelegs and ThC roll for the middle and hind legs.

Each DoF was controlled by two coupled CPGs that drove the extensor and antagonistic flexor muscles. We assumed left-right body symmetry and optimized intraleg joint phase differences and muscle parameters for the right legs, mirroring these results for the left legs. In the same manner, we optimized the phase differences between the coxae flexor CPGs and mirrored them for the coxae extensor CPGs. Thus, we could connect 36 coupled oscillators in a minimal configuration to remove redundancy and reduce the optimization search space (Figure 6*A*). Finally, to permit a wide range of joint movements, each CPG’s intrinsic frequency was set as an open parameter, whose limits were constrained to biologically relevant frequencies observed from real fly joint movements during walking [28, 65](Figure S13). In total, 63 open parameters were optimized including CPG intrinsic frequencies, CPG phase differences, and muscle parameters (see Materials and Methods).

We performed multi-objective optimization [66] using the NSGA-II genetic algorithm [67] to identify neuromuscular parameters that drove walking gaits satisfying two high-level objective functions: forward speed and static stability. Notably, these objectives can be inversely correlated: fast walking might be achieved by minimizing stance duration and reducing static stability. Forward speed was defined as the number of backward ball rotations within a fixed period of time and quantified as fictive distance traveled (Figure 6*B*, top). Static stability refers to the stability of an animal’s given pose if, hypothetically, tested while immobile. This metric can be quantified during walking as the minimal distance between the model’s center-of-mass (COM) and the closest edge of the support polygon formed by the legs in stance phase (i.e., in contact with the ground). This means that the closer the COM is to the center of the support polygon, the higher the static stability score. (Figure 6*B*, bottom). Additionally, we defined four penalties to discourage unrealistic solutions including those with excessive joint velocities (these cause jittering or muscle instability), speeds slower or faster than real locomotion (a ‘moving boundary’), as well as joint angle ranges of motion and duty factors that violate those observed in real flies. Because the optimizer minimizes the objective functions, we inverted the sign for both functions. Thus, during optimization the Pareto front of best solutions evolved toward more negative values (Figure S14*A*) and forward walking speeds became faster over generations (Figure 6*C*)**(Video 9)**.

To more deeply investigate our optimization results, we examined three individual solutions from the final generation. These were: (i) the fastest solution, (ii) the most stable solution, and (iii) a ‘trade-off’ solution that was the best compromise between speed and static stability (see Methods for a precise mathematical definition) (Figure 6*D*). By generating gait diagrams for each of these solutions, we found a diversity of strategies−non-tripod gaits were observed in all generations (Figure S14*B*) even after objectives were maximized and penalties minimized at generation 60 (Figure S14*C*). However, the trade-off solution−a compromise between speed and static stability−closely resembled a typical insect tripod gait [28, 68], supporting the notion that tripod locomotion satisfies a need for stability during fast insect walking [24].

Because NeuroMechFly provides access to neuromuscular dynamics and physical interactions, we could also analyze then further analyze how these underlying quantities give rise to optimized locomotor gaits. To illustrate this, we focused on the femur flexors of each leg for the ‘trade-off’ solution (Figure 6*F*). As expected for a tripod gait, stance and swing phases of the left front (LF) and hind (LH) legs were coordinated with those of the right middle (RM) leg. This coordination implies that the middle and hind legs CPG activities (Figure 6*F*, top, green and brown) are in phase with each other and phase shifted by 180º with respect to the front leg (Figure 6*F*, top, orange). This is because, during stance phases, the front legs flex while the middle and hind legs extend. However, for the tripod generated by other three legs, the CPG activity of the left, middle (LM) femur was phase shifted with respect to the right front (RF) and hind (RH) legs (Figure 6*F*, top, red). Torques were highest for the hind legs, suggesting an important role for driving ball rotations (Figure 6*F*, middle, purple and brown). Finally, we confirmed that the increased torque of the hind legs was associated with a larger range of motion as measured by joint angles (Figure 6*F*, bottom).

These results illustrate how, by combining our biomechanical exoskeleton with neuromuscular elements and an optimization framework, we could discover control strategies that maximize highlevel behavioral objectives and minimize penalties informed by real measurements of *Drosophila*. For these solutions, neuromuscular dynamics, collisions, and forces could then be further examined because of their instantiation within a physics-based simulation environment.

## 3 Discussion

Here we have introduced NeuroMechFly, a computational model of adult *Drosophila* that can be used for biomechanical, and−by also including available neural and muscle models−neuromechanical studies. We first illustrated a biomechanical use case in which one can estimate joint torques and contact forces including ground-reaction forces and body part collisions by replaying real, measured fly walking and grooming. In the future, directly through force measurements [69, 70] or indirectly through recordings of proprioceptive and tactile neurons [38, 71], these estimates might be further validated. Next, we demonstrated a neuromechanical use case by showing how high-level optimization of a neural network and muscles could be used to discover and more deeply study locomotor controllers. Although here we optimized for speed and static stability during tethered locomotion, NeuroMechFly can also locomote without body support, opening up the possibility of optimizing neuromuscular controllers for diverse, untethered behaviors.

### 3.1 Limitations and future extensions of the biomechanical module

The biomechanical exoskeleton of NeuroMechFly can benefit from several near-term extensions by the community. First, actuation is currently only implemented for leg joints. Additional effort will be required to actuate other body parts including the head, or abdomen by defining their DoFs, joint angle ranges and velocities based on 3D pose measurements. Second, the model currently achieves compliant joints during kinematic replay through position control (akin to a spring-and-damper) in PyBullet. However, future work may include implementing compliant joints with stiffness and damping based on measurements from real flies. Third, NeuroMechFly employs rigid bodies that do not reflect the flexibility of insect cuticle. Although our modeling framework could potentially include soft-bodied elements−these are supported by the underlying physics engine−we have chosen not to because it would first require challenging measurements of cuticular responses to mechanical stresses and strains (i.e. Young’s modulus) [72,73], and this would increase the model’s computational complexity, making it less amenable to evolutionary optimization. NeuroMechFly currently supports flexibility in terms of compliance because the muscle model includes stiffness and damping terms. Additionally, the fact that kinematic replay is already accurate−with similar real and simulated joint angle and end-effector positions−suggests that modeling additional cuticular deformations might only have negligible effects. Therefore, we currently offer what we believe to be a practical balance between accuracy and computational cost. Finally, future iterations of our biomechanical model might also include forces that are observed at small scales, including Van der Waals and attractive capillary forces of footpad hairs [74].

### 3.2 Limitations and future extensions of the neuromuscular modules

In addition to its biomechanical exoskeleton, NeuroMechFly includes modules for neural controllers, muscle models, and the physical environment (Figure 7). These interact with one another to generate rich *in silico* motor behaviors. Each of these modules can be independently modified in future work to improve biological interpretability, computational efficiency, and increase the range of possible experiments. First, more detailed neural controllers could already be implemented including Integrate-and-Fire, or Hodgkin-Huxley type neurons [15]. This would aid in the comparison of discovered artificial neural networks and their dynamics with measured connectomes [40, 41] and functional recordings [38], respectively. Second, to increase the realism of movement control, Hill-type muscle models that have nonlinear force generation properties could be implemented based on species-specific muscle properties−slack tendon lengths, attachment points, maximum forces, and pennation angles [58, 75]. Third, to study more complex motor tasks, one can already use the PyBullet framework [42] to increase the complexity of the physical environment. For example, one can study locomotor stability by introducing external objects **(Video 10)**, or locomotor strategies for navigating heightfield terrains.

**Figure 7:**
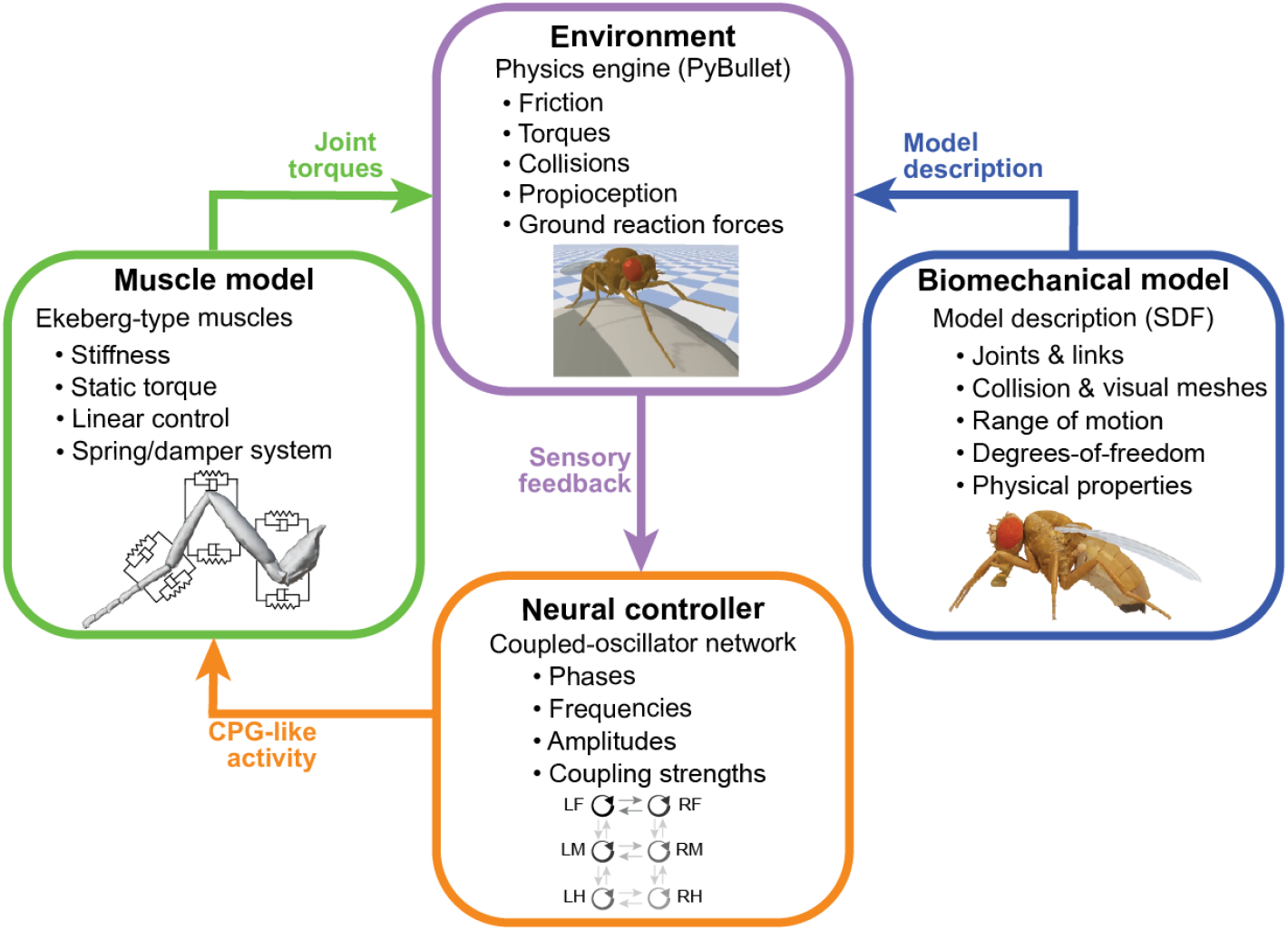
Modules that can be independently modified in NeuroMechFly. A neural controller’s output drives muscles to move a biomechanical model in a physics-based environment. Each of these modules can be independently modified or replaced within the NeuroMechFly simulation framework. The controller generates neural-like activity to drive muscles. These muscles produce torques to operate a biomechanical model embedded in PyBullet’s physics-based environment. When replacing any module it is only necessary to preserve the inputs and outputs (colored arrows).

In the near-term, we envision that NeuroMechFly will be used to test theories for neuromechanical behavioral control. For example, one might investigate the respective roles of feedforward versus feedback mechanisms in movement control (i.e., to what extent movements are generated by central versus sensory-driven signals). This can be tested by systematically modifying coupling strengths and sensory feedback gains in the simulation. Outcomes may then be experimentally validated. In the longer-term, this modeling framework might also be used in closed-loop with ongoing neural and behavioral measurements. Real-time 3D poses might be replayed through NeuroMechFly to predict joint torques and contact forces. These leg state predictions might then inform the delivery of perturbations to study how proprioceptive or tactile feedback are used to achieve robust movement control. In summary, NeuroMechFly promises to accelerate the investigation of how passive biomechanics and active neuromuscular control orchestrate animal behavior, and can serve as a bridge linking fundamental biological discoveries to applications in artificial intelligence and robotics.

## 4 Materials and Methods

### 4.1 Constructing an adult *Drosophila* biomechanical model

#### 4.1.1 Preparing adult flies for x-ray microtomography

The protocol used to prepare flies for microtomography was designed to avoid distorting the exoskeleton. We observed that traditional approaches for preparing insects for either archival purposes or for high resolution microscopy, including scanning electron microscopy [76], result in the partial collapse or bending of some leg segments and dents in the exoskeleton of the thorax and abdomen. These alterations mostly occur during the drying phase and while removal of ethanol by using supercritical carbon dioxide drying reduces these somewhat, it is still not satisfactory. We therefore removed this step altogether, and instead embedded flies in a transparent resin. This resulted in only a small surface artifact over the dorsal abdominal segments A1, A2, and A3.

Flies were heavily anaesthetized with CO_2_ gas, then carefully immersed in a solution of 2% paraformaldehyde in phosphate buffer (0.1M, pH 7.4) containing 0.1% Triton 100, to ensure fixative penetration, and left for 24 h at 4°C. Care was taken to ensure the flies did not float on the surface, but remained just below the meniscus. They were then washed in 0.1M cacodylate buffer (2 × 3 min washes), and placed in 1% osmium tetroxide in 0.1M cacodylate buffer, and left at 4°C for an additional 24 h. Flies were then washed in distilled water and dehydrated in 70% ethanol for 48 h, followed by 100% ethanol for 72 h, before being infiltrated with 100% LR White acrylic resin (Electron Microscopy Sciences, US) for 24 h at room temperature. This was polymerised for 24 h at 60°C inside a closed gelatin capsule (size 1; Electron Microscopy Sciences) half-filled with previously hardened resin to ensure the insect was situated in the center of the final resin block, and away from the side.

#### 4.1.2 X-ray microtomography

We glued the sample onto a small carbon pillar and scanned it using a 160 kV open type, microfocus X-ray source (L10711/-01; Hamamatsu Photonics K.K., Japan). The X-ray voltage was set to 40 kV and the current was set to 112 uA. The voxel size was 0.00327683 mm. To perform the reconstruction, we used X-Act software from the microtomography system developer (RX-solutions, Chavanod, France) obtaining a stack of 982 tiff images of 1046×1636 pixels each.

#### 4.1.3 Building a polygonal mesh volume from processed microtomography data

First, we isolated cuticle and wings from the microtomography data using Fiji [77]. We selected 360 images from the tiff stack as the region of interest (ROI) beginning at slice 300. The tiff stack with the ROI was then duplicated. The first copy was binarized using a threshold value of 64 to isolate the cuticle. The second copy was cropped to keep the upper half of the image−where the wings are− and then binarized using a lower threshold value of 58. Finally, we applied a closing morphological operation to isolate the wings. Both binarized stacks were stored as tiff files.

We developed custom Python code to read the tiff stacks, and to fill empty holes within the body and wings. Finally, we used the Lewiner marching cubes algorithm [43] (implemented in the scikitimage package [78]) to obtain a polygon mesh for each stack. Both meshes were then exported to a standard compressed mesh storage format.

#### 4.1.4 Separating and reassembling articulated body parts

We used Blender (Foundation version 2.81 [79]) to clean and manipulate polygon meshes obtained from microtomography data.

After importing these meshes into Blender, we removed noise by selecting all vertices linked to the main body (or wings), inverting the selection, and deleting these vertices. We explored the resulting meshes, looking for spurious features, and then manually selected and deleted the related vertices. We obtained 65 body segments (Table 1) based on [80]. More recent literature corroborated these propositions for body morphology and joint degrees-of-freedom. We manually selected and deleted vertices from our imported 3D body and wing models. Segments were then separated at joint locations based on published morphological studies. We made some simplifications. Most notably, in the antennae, we considered only one segment instead of three because cutting this small element into a few pieces would alter its morphology.

Each wing was separated into an individual segment from the wing model. The body model was separated into 63 segments as described below. The abdomen was divided into five segments according to tergite divisions. The first and second tergites were combined as the first segment (A1A2), and the last segment (A6) included the sixth to tenth tergites. Each antenna was considered a single segment and separated from the head capsule at the antennal foramen. Both eyes and the proboscis were separated from the head. The latter was divided into two parts, the first containing the rostrum (Rostrum), and the second containing the haustellum and labellum (Haustellum). Each leg was divided in eight parts: the coxa, trochanter/femur, tibia, and five tarsal segments. The thorax was considered a single segment and only the halteres were separated from it.

Each segment was processed in Blender to obtain closed meshes. First, a remesh modifier was used in ‘smooth mode’, with an octree depth of 8, and a scale of 0.9 to close the gaps generated in the meshes after been separated from the original model. Smooth shading was enabled and all disconnected pieces were removed. Then, we used ‘sculpt mode’ to manually compensate for depressions/collapses resulting from the microtomography preparation, or from separating body segments.

Then, all segments were copied into a single *.blend file and rearranged into a natural resting pose (Figure 2F). We made the model symmetric to avoid inertial differences between contralateral legs and body parts. For this, we used the more detailed microtomography data containing the right side of the fly. First, the model was split along the longitudinal plane using the bisect tool. Then the left side was eliminated and the right side was duplicated and mirrored. Finally, the mirrored half was repositioned as the left side of the model, and both sides of the head capsule, rostrum, haustellum, thorax, and abdominal segments were joined.

At this point, the model consisted of approximately nine million vertices, an intractable number for commonly used simulators. We therefore used the decimate tool to simplify the mesh and collapse its edges at a ratio of 1% for every segment. This resulted in a model with 87,000 vertices that conserved the most important details but eliminated small bristles and cuticular textures.

#### 4.1.5 Rigging the Blender model

We added an Armature object alongside our model to build the skeleton of the fly. To actuate the model, we created a ‘bone’−a tool in Blender that is used to animate characters−for each segment. Bones were created such that the thorax would be the root of the skeleton and each bone would be the child of its proximal bone, as indicated in Table 1. Then, the bones were positioned along the longitudinal axis of each segment with their heads and tails over the proximal and distal joints, respectively. Each joint was positioned at a location between neighboring segments. Each bone inherited the name of its corresponding mesh.

We used the Custom Properties feature in Blender to modify the properties of each bone. These properties can be used later in a simulator to e.g., define the maximum velocity, or maximum effort of each link. Furthermore, we added a limit rotation constraint (range of motion) to each axis of rotation (DoF) for every bone. The range of motion for each rotation axis per joint was defined as −180^*°*^ to 180^*°*^ to achieve more biorealistic movements. Because, to the best of our knowledge, there are no reported angles for these variables, these ranges of motion should be further refined once relevant data become available. The DoF of each bone (segment) were based on previous studies [44, 81, 82] (see Table 1). Any bone can be rotated in Blender to observe the constraints imposed upon each axis of rotation. These axes are defined locally for each bone.

Finally, we defined a ‘zero-position’ for our model. Most bones were positioned in the direction of an axis of rotation (Figure S7). Each leg segment and the proboscis were positioned along the *Z* axis. Each abdominal segment and the labellum were positioned along the *X* axis. Wings, eyes, and halteres were positioned along the *Y* axis. The head and the antennae are the only bones not along a rotational axis: the head is rotated 20^*°*^ along the *Y* axis, and the antennae are rotated 90^*°*^ with respect to the head bone. Positioning the bones along axes of rotation makes it easier to intuit a segment’s position with its angular information and also more effectively standardizes the direction of movements.

#### 4.1.6 Exporting the Blender model into the Bullet simulation engine

We used a custom Python script in Blender to obtain the name, location, global rotation axis, range of motion, and custom properties for each bone. As mentioned above, the axes of rotation are defined locally for each bone. Therefore, our code also transforms this information from a local to a global reference system, obtaining the rotation matrix for each bone.

We used the Simulation Description Format (SDF, http://sdformat.org/) convention to store the model’s information. This format consists of an *.xml file that describes objects and environments in terms of their visualization and control. The SDF file contains all of the information related to the joints (rotational axes, limits, and hierarchical relations) and segments (location, orientation, and corresponding paths of the meshes) of the biomechanical model. We can modify this file to add or remove segments, joints, or to modify features of existing segments and joints. To implement joint DoFs, we used hinge-type joints because they offer more freedom to control individual rotations. Therefore, for joints with more than one DoF, we positioned in a single location as many rotational joints as DoFs needed to describe its movement. The parenting hierarchy among these extra joints was defined as roll-pitch-yaw. The mass and collision mesh were related to the segment attached to the pitch joint−present in every joint of the model. The extra segments were defined with a zero mass and no collision shape.

Our model is based upon the physical properties of a real fly. The full body length and mass of the model are set to 2.8 *mm* and 1 *mg*, respectively. To make the center of mass and the rigid-body dynamics of the model more similar to a real fly, rather than having a homogeneous mass distribution, we used different masses (densities) for certain body parts as measured in a previous study [83]. Specifically, these masses were: head (0.125 *mg*), thorax (0.31 *mg*), abdomen (0.45 *mg*), wings (0.005 *mg*), and legs (0.11 *mg*).

In PyBullet, contacts are modeled based on penetration depth between any two interacting bodies. The contact parameters are set to 0.02 units of length (1 unit = 1 m in SI units). It is preferable to have the bodies of size larger than 0.02 units. Therefore, we performed dynamic scaling to rescale the model, the physical units, and quantities such as gravity while preserving the dynamics and improving the numerical stability of the model. Notably, we are not compromising the dynamics of the simulated behaviors. Specifically, we scaled up the units of mass and length when setting up the physics of the simulation environment, and then scaled down the calculated values when recording the results. Therefore, the physics engine was able to compute the physical quantities without numerical errors, and the model could also more accurately reflect the physics of a real fly.

#### 4.1.7 Comparing leg sizes between NeuroMechFly and real flies

We dissected the right legs from ten wild-type female adult flies, 2-4 days-post-eclosion. Flies were cold anesthetized using ice. Then the legs were removed using forceps from the sternal cuticle to avoid damaging the coxae. Dissected legs were straightened onto a glass slide and fixed with UV-curable glue (Figure S1A). We used a Leica M205 C stereo microscope to take images from the legs placed next to a 0.5 mm graduated ruler. Joints in the legs were manually annotated and then distances between them were measured in pixels and converted to mm using the ruler as a reference. Lengths between joints were compared to rigged bone lengths in NeuroMechFly.

### 4.2 Kinematic replay and analysis

#### 4.2.1 Forward walking data

We recorded spontaneous behaviors from wild-type females 3-4 days-post-eclosion. Flies were mounted on a custom stage and allowed to acclimate for 15 min on an air-supported spherical treadmill [38]. Experiments were conducted in the evening Zeitgeber time. Flies were recorded five times for 30 s at 5 min intervals. Data were excluded if forward walking wasn’t present for at least five continuous seconds in 10 s windows. To record data, we used a 7-camera system as in [33]. However, we replaced the front camera’s InfiniStix lens with a Computar MLM3X-MP lens at 0.3x zoom to visualize the spherical treadmill. After the fifth trial of each experiment, we recorded an extra 10 s trial, having replaced the lens from a lateral camera with another Computar MLM3X-MP lens. We used these images to calculate the longitudinal position of the spherical treadmill with respect to the fly for the preceding five trials.

#### 4.2.2 Foreleg/antennal grooming data

Data for kinematic replay of foreleg/antennal grooming were obtained from a previous study describing DeepFly3D, a deep learning-based 3D pose estimation tool [33]. These data consist of images from seven synchronized cameras obtained at 100 fps (https://dataverse.harvard.edu/dataverse/DeepFly3D). Time axes (Figure 5*E, F*) correspond to time points from the original, published videos. Data were specifically obtained from experiment #3, taken of an animal (#6) expressing aDN-GAL4 driving UAS-CsChrimson.

#### 4.2.3 Processing 3D pose data

We used DeepFly3D v0.4 [33] to obtain 3D poses from the images acquired for each behavior. 2D poses were examined using the GUI to manually correct 10 frames during walking and 72 frames during grooming. DeepFly3D, like many other pose estimation softwares, uses a local reference system based on the cameras’ positions to define the animal’s pose. Therefore, we first defined a global reference system for NeuroMechFly from which we could compare data from experiments on different animals (see Figure S7).

Aligning both reference systems consisted of six steps. First, we defined the mean position of each Thorax-Coxa (ThC) keypoint as fixed joint locations. Second, we calculated the orientation of the vectors formed between the hind and middle coxae on each side of the fly with respect to the global x-axis along the dorsal plane. Third, we treated each leg segment independently and defined its origin as the position of the proximal joint. Fourth, we rotated all data points on each leg according to its side (i.e., left or right) and previously obtained orientations. Fifth, we scaled the real fly’s leg lengths for each experiment to fit NeuroMechFly’s leg size: A scaling factor was calculated for each leg segment as the ratio between its mean length throughout the experiment and the template’s segment length and then each data point was scaled using this factor. Finally, we used the NeuroMechFly exoskeleton as a template to position all coxae within our global reference system; the exoskeleton has global location information for each joint. Next, we translated each data point for each leg (i.e. CTr, FTi, and TiTa joints) with respect to the ThC position based on this template.

#### 4.2.4 Calculating joint angles from 3D poses

We considered each leg a kinematic chain and calculated the angle of each DoF to reproduce real poses in NeuroMechFly. We refer to this process as ‘kinematic replay’. Angles were obtained by computing the dot product between two vectors with a common origin. We obtained 42 angles in total, seven per leg. The angles’ names correspond to the rotational axis of the movement−roll, pitch, or yaw−for rotations around the anterior-posterior, mediolateral, and dorsoventral axes, respectively.

The thorax-coxa joint (ThC) has three DoFs. The yaw angle is measured between the dorsoventral axis and the coxa’s projection in the transverse plane. The pitch angle is measured between the dorsoventral axis and the coxa’s projection in the sagittal plane. To calculate the roll angle, we aligned the coxa to the dorsoventral axis by rotating the kinematic chain from the thorax to the FTi joint using the yaw and pitch angles. Then we measured the angle between the anterior-posterior axis and the projection of the rotated FTi in the dorsal plane.

Initially, we considered only a pitch DoF for the CTr joint. This was measured between the coxa and femur’s longitudinal axis. Subsequently, we discovered that a CTr roll DoF would be required to accurately match the kinematic chain. To calculate this angle, we rotated the tibia-tarsus joint (TiTa) using the inverse angles from the coxa and femur and measured the angle between the anterior-posterior axis and the projection of the rotated TiTa in the dorsal plane.

The pitch angle for the FTi was measured between the femur and tibia’s longitudinal axis. The pitch angle for the TiTa was measured between the tibia and tarsus’s longitudinal axis. The direction of rotation was calculated by the determinant between the vectors forming the angle and its rotational axis. If the determinant was negative, the angle was inverted.

To demonstrate that the base six DoFs were not sufficient for accurate kinematic replay, we also compared these results to angles obtained using inverse kinematics. In other words, we assessed whether an optimizer could find a set of angles that could precisely match our kinematic chain using only these six DoFs. To compute inverse kinematics for each leg, we used the optimization method implemented in the Python IKPy package (L-BFGS-B from Scipy). We defined the zero-pose as a kinematic chain and used the angles from the first frame as an initial position (seed) for the optimizer.

#### 4.2.5 Calculating forward kinematics and errors with respect to 3D poses

To quantify the contribution of each DoF to kinematic replay, we used the forward kinematics method to compare original and reconstructed poses. Since 3D pose estimation noise causes leg segment lengths to vary, we first fixed the length of each segment as its mean length across all video frames.

We then calculated joint angles from 3D pose estimates with the addition of each DoF (see previous section). We formed a new kinematic chain including the new DoF. This kinematic chain allowed us to compute forward kinematics from joint angles, which were then compared with 3D pose estimates to calculate an error. We performed an exhaustive search to find angles that minimize the overall distance between each 3D pose joint position and that joint’s position as reconstructed using forward kinematics. The search spanned from −90^*°*^ to 90^*°*^ with respect to the ‘zero pose’ in 0.5^*°*^ increments. The error between 3D pose-based and angle-based joint positions per leg was calculated as the average distance across every joint. We note that differences in errors can vary across legs and leg pairs because each joint’s 3D pose estimate is independent and each leg acts as an independent kinematic chain adopting its own pose. Thus, errors may also be asymmetric across the body halves. As well, errors integrate along the leg when using forward kinematics (FK) for walking (Figure S2) and for grooming (Figure S3). By contrast, inverse kinematics (IK) acts as an optimizer and minimizes the error at the end of the kinematic chain (i.e., where the FK error is highest) for walking (Figure S2D) and for grooming (Figure S3D). This explains why errors using FK are generally higher than those using IK−with the exception of adding a roll degree-of-freedom at the Coxa-Trochanter joint. To normalize the error with respect to body length, we measured the distance between the antennae and genitals in our Blender model (2.88 *mm*). Errors were computed using 400 frames of data: frames 300-699 for forward walking from fly 1 and frames 0-399 for foreleg/antennal grooming.

We ran a Kruskal-Wallis statistical test to compare kinematic errors across the eight methods used. We then applied a posthoc Conover’s test to perform a pairwise comparison. We used the Holm method to control for multiple comparisons. The resulting p-value matrices for walking and foreleg/antennal grooming behaviors are shown in Table 2 and Table 3, respectively. Our statistical tests suggested that adding a CTr roll DoF uniquely improved kinematic replay compared with all other methods.

#### 4.2.6 Transferring real 3D poses into the NeuroMechFly reference frame

To incorporate the additional CTr roll DoF into NeuroMechFly, we enabled rotations along the *z* axis of CTr joints. Then, we created new SDF configuration files using custom Python scripts to include a CTr roll DoF for each leg. To simulate the fly tethering stage used in our experiments, we added three support joints (one per axis of movement) that would hold our model in place. We removed these supports for ground walking experiments **(Videos 8 and 10)**.

We used position control for each joint in the model. We fixed the position of non-actuated joints to the values shown in Table 4. The actuated joints (i.e. the leg joints) were controlled to achieve the angles calculated from 3D pose data. The simulation was run with a time step of 0.5 *ms*, allowing PyBullet to accurately perform numerical calculations. Since the fly recordings were only captured at 100 fps, we up-sampled and interpolated pose estimates to match the simulation time steps before calculating joint angles.

#### 4.2.7 Comparing real and simulated spherical treadmill rotations

We obtained spherical treadmill rotational velocities from real experiments using Fictrac [57]. We also obtained the relative inclination of each tethered fly (ϕ) (Figure S9A) as the angle between the ground plane and the axis between the hind leg ThC joint and the dorsal part of the neck. Finally, we estimated the position of the ball with respect to the fly from both front and lateral views (Figure S9B-C) by identifying the ball and fly using a Hough transform and standard thresholding, respectively. For axes observed from both views, we averaged the expected position.

For the simulated environment we created a spherical body in PyBullet with three hinge joints along the *x, y*, and *z* axes, allowing our sphere to rotate in each direction like a real spherical treadmill. Rolling and spinning frictions were set to zero to obtain virtually frictionless conditions similar to a real treadmill floating on air. The mass of the simulated spherical treadmill was set to 54.6 *mg*: the measured mass of the real foam sphere. Finally, the sphere’s diameter was measured and set into the simulation as 9.96 *mm*.

We ran kinematic replay of walking by setting the simulated spherical treadmill position and fly inclination based on measurements from experimental images. We used predefined values for kinematic replay of grooming. Then, we empirically determined the following parameters:

- Global ERP = 0.0
- Friction ERP = 0.0
- Solver iterations = 1000
- Treadmill lateral friction = 1.3

After running the simulation, we compared the rotational velocities estimated for each axis with the real velocities obtained with Fictrac. First, we smoothed both Fictrac and estimated signals using a median filter with a window size of 0.1 s. Second, we interpolated Fictrac data from time steps of 0.1 s (100 fps) to the simulation time step. Then, we established each signal’s baseline as the mean of the first 0.2 s of data. Finally, we computed the Spearman correlation coefficient (*ρ*) to assess correlations of forward, lateral, and heading (yaw) velocities for both signals.

#### 4.2.8 Constraint parameter sensitivity analysis

Simulated spherical treadmill velocity estimates depend on constraint force mixing (CFM) and contact error reduction (contact ERP) parameters. These parameters change the ‘softness’ of joint and contact constraints in the physics engine. Therefore we performed a sensitivity analysis to determine the best combination of CFM and ERP. CFM values were swept from 0 to 10, and ERP from 0 to 1.0. Then, we ran a simulation for each of 121 combinations. We assessed their performance by calculating the Spearman correlation coefficient for each axis (Figure S8A-C).

Finally, to select optimal parameter values, we applied a weighted sum to the results as shown in Equation 1:

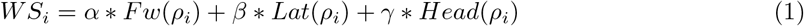

where Fw, Lat, and Head are the rotational axes, *ρ*_*i*_ is the Spearman correlation coefficient obtained for each CFM-ERP combination, and *α, β*, and *γ* are the standard deviation contributions for each axis calculated as shown in Equations 2, 3, and 4, respectively. Therefore, we favored the axis with the largest amplitude of variation.

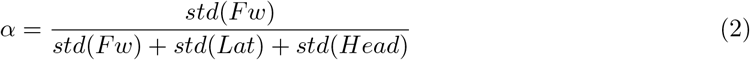

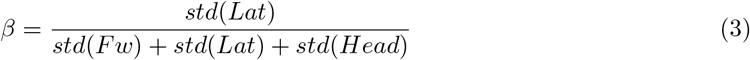

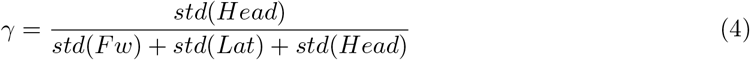

Finally, we normalized WS (NWS) with respect to its maximum and minimum values (Figure S8D). Consequently, a combination with NWS equal to 1 was selected: CFM = 3 and ERP = 0.1.

#### 4.2.9 Controller gain sensitivity analysis

We performed kinematic replay using a built-in PD position controller in PyBullet [42]. A PD controller was used rather than the more widely known PID controller because the integral component (‘I’ in PID) is mainly used to correct steady state errors (e.g., while maintaining a fixed posture). Thus, it is not used for time-varying postures like those during locomotion. We used PyBullet’s built-in position control method because it operates with proportional and derivative gains that are stable and efficient. This PD controller minimizes the error:

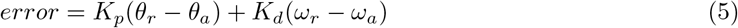

where *θ*_*r*_ and *θ*_*a*_ denote reference and actual positions, *ω*_*r*_ and *ω*_*a*_ are desired and actual velocities, and *K*_*p*_ and *K*_*d*_ are proportional and derivative gains, respectively, which provides some compliance in the model.

Because the outputs of our model−dynamics of motion−depend on the controller gains *K*_*p*_ and *K*_*d*_, we first systematically searched for optimal gain values. To do this, we ran the simulation’s kinematic replay for numerous *K*_*p*_ and *K*_*d*_ pairs, ranging from 0.1 to 1.0 with a step size of 0.1 (i.e., 100 simulations in total). Target position and velocity signals for the controller were set as the calculated joint angles and angular velocities, respectively. To compute joint angular velocities, we used a Savitzky–Golay filter with a first-order derivative and a time-step of 0.5 ms on the joint angles. Feeding the controller with only the joint angles could also achieve the desired movements of the model. However, including the velocity signal ensured that the joint angular velocities of the fly and the simulation were properly matched. We then calculated the mean squared error (MSE) between the ground truth−joint angles obtained by running our kinematic replay pipeline on pose estimates from DeepFly3D [33]−and joint angles obtained from PyBullet. Then, we averaged the MSE values across the joints in one leg, and summed the mean MSEs from each of six legs to obtain a total error. We made the same calculations for the joint angular velocities as well. Our results (Figure S4) show that our biomechanical model can replicate real 3D poses while also closely matching real measured velocities. In particular, an MSE of 360 (*rad/sec*)^2^ for the six legs corresponds approximately to 7.74 *rad/sec* per leg, i.e., 1.27 *Hz*. This is acceptable given the rapid, nearly 20 Hz, leg movements of the real fly.

After validating the accuracy of kinematic replay, we performed a sensitivity analysis to measure the impact of varying controller gains on the estimated torques and ground reaction forces. This analysis showed that torques and ground reaction forces are highly sensitive to changing proportional gains (*K*_*p*_) (Figure S5) but are robust to variations in derivative gain (*K*_*d*_). These results are expected since high proportional gains cause “stiffness” in the system whereas derivative gains affect the “damping” in a system’s response. We observed rapid changes in estimated torques and ground reaction forces at high *K*_*p*_ values (Figure S5). Notably, in principle there can also be internal forces affecting contact forces. For example, a fly’s legs can squeeze the spherical treadmill with different internal forces but have identical postures.

As shown in Figure S4, our model can match the real kinematics closely for almost every controller gain combination except for the low *K*_*p*_, *K*_*d*_ band. By contrast, varying the gains proportionally increased the torque and force readings. Because there are no experimental data to validate these physical quantities, we selected gain values corresponding to intermediate joint torques and ground contact forces (Figure S5). Specifically, we chose 0.4 and 0.9 for *K*_*p*_ and *K*_*d*_, respectively. These values were high enough to generate smooth movements, and low enough to reduce movement stiffness.

#### 4.2.10 Comparing tethered and flat ground walking

To test the ability to run NeuroMechFly in an untethered context, we replayed the kinematics of a tethered walking experiment (Figure 4) but removed body supports and placed the model on the floor. To remove body supports, we deleted the corresponding links from the model’s description (SDF configuration file). The physics engine parameters remained the same. The lateral friction for the floor was set to 0.1.

#### 4.2.11 Application of external perturbations

To test the stability of the untethered model walking over flat ground, we set the floor’s lateral friction to 0.5 and introduced external perturbations. Specifically, we propelled solid spheres at the model according to the following equation of motion,

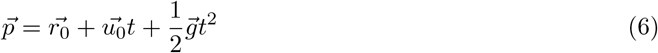

where, 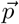 is the 3D target position(fly’s center of mass), 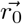 is the initial 3D position of the sphere, 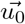 is the initial velocity vector, 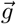 is the external acceleration vector due to gravity in the z-direction, *t* is the time taken by the sphere to reach the target position 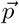 from 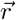 with an initial velocity 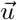. The mass of the sphere was 3 *mg* and its radius 50 *µm*. Spheres were placed at a distance of 2 *mm* from the fly’s center of mass in the y-direction. With *t* set to 20 *ms*, the initial velocity of the projectile was computed using Equation 6. The spheres were propelled at the model every 0.5 *s*. Finally, at 3 *s* into the simulation, a 3 *g* sphere with a radius of 150 *µm* was propelled at the fly to topple it over **(Video 10)**.

#### 4.2.12 Analyzing NeuroMechFly’s contact and collision data

The PyBullet physics engine generates forward dynamics simulations and collision detections. We plotted joint torques as calculated from PyBullet. To infer ground reaction forces (GRFs), we computed and summed the magnitude of normal forces resulting from contact of each tarsal segment with the ball. Gait diagrams were generated by thresholding GRFs; a leg was considered to be in stance phase if its GRFs was greater than zero. These gait diagrams were compared with a ground truth (Figure S10) obtained by manually annotating when the legs were in contact with the ball for each video frame. Gait prediction accuracy was calculated by dividing the frames correctly predicted as being in stance or swing over the total number of frames.

Self-collisions are disabled by default in PyBullet. Therefore, for kinematic replay of grooming, we enabled self-collisions between the tibia and tarsal leg segments, as well as the antennae. We recorded normal forces generated by collisions between (i) the right and left front leg, (ii) the left front leg and left antenna, and (iii) the right front leg and right antenna. Grooming diagrams were calculated as for gait diagrams: a segment experienced a contact/collision if it reported a normal force greater than zero.

#### 4.2.13 Comparing grooming behaviors as a function of NeuroMechFly’s morphological accuracy

We replayed foreleg/antennal grooming kinematics (Figure 5) for three conditions to assess the degree to which biomechanical realism is important for collision estimation. We tested two experimental conditions: one in which both front legs were modelled as sticks, and one in which the front legs as well as the antennae were modelled as sticks. Notably, multisegmented tarsi are not found in other published insect stick models [64]. Thus, as for our previous model [24], each stick leg consisted of four segments: coxa, trochanter/femur, tibia, and one tarsal segment. Each leg and antennal stick segment had a diameter equal to the average diameter of the corresponding segment in our more detailed NeuroMechFly model. These changes were accomplished by modifying the model’s description (SDF configuration file) and by changing the collision and visual attributes for each segment of interest.

### 4.3 Neural network parameter optimization

#### 4.3.1 CPG network architecture

For evolutionary optimization of neuromusculuar parameters, we designed a CPG-based controller composed of 36 nonlinear oscillators (Figure 6), as for a previous investigation of salamander locomotion [62]. These CPGs consisted of mathematical oscillators that represent neuronal ensembles firing rhythmically in the Ventral Nerve Cord (VNC) [84]. The CPG model was governed by the following system of differential equations:

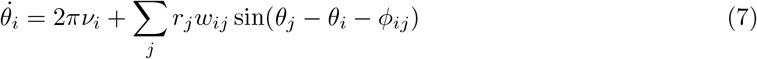

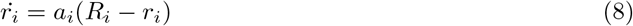

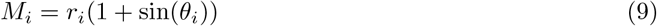

where the state variables−phase and amplitude of the oscillator *i*−are denoted *θ*_*i*_ and *r*_*i*_, respectively; *ν*_*i*_ and *R*_*i*_ represent oscillator *i*’s intrinsic frequency and amplitude, *a*_*i*_ is a constant. The coupling strength and phase bias between the oscillator *i* and *j* are denoted *w*_*ij*_ and *ϕ*_*ij*_, respectively.

During optimization, for the entire network of coupled oscillators, we set the intrinsic frequency *v* as an open parameter ranging from 6 to 10 Hz, matching the frequencies of our measured *Drosophila* joint angle movements and reported stepping frequencies [65]. The intrinsic amplitude *R* was set to 1, and the constant *a*_*i*_ was set to 25. To ensure a faster convergence to a phase-locked regime between oscillators, we set coupling strengths to 1000 [85]. *M*_*i*_ represents the cyclical activity pattern of neural ensembles activating muscles. We solved this system of differential equations using the explicit Runge-Kutta method of 5th-order with a time step of 0.1 ms.

Each oscillator pair sends cyclical bursts to flexor and extensor muscles which apply antagonistic torques to the corresponding revolute joint. We considered three DoFs per leg that were sufficient for locomotion in previous hexapod models [64] and that had the most pronounced joint angles (Figure S13). These DoFs were (i) ThC pitch for the front legs, (ii) ThC roll for the middle and hind legs, and (iii) CTr pitch and FTi pitch for all legs. Thus, there were three pairs of oscillators optimized per leg, for a total of 36. We coupled (i) the intraleg oscillators in a proximal to distal chain, (ii) the interleg oscillators in a tripod-like fashion (the ipsilateral front and hind legs to the contralateral middle leg from anterior to posterior), (iii) both front legs to each other, and (iv) coxa extensor and flexor oscillators to one another. Intraleg coordination is equally important to generate a fly-like gaits since stance and swing phases depend on intrasegmental phase relationships. For this reason, both interleg (phase relationships between ThC joints) and intraleg (phase relationships within each leg) couplings were optimized for one half of the body and mirrored on the other.

#### 4.3.2 Muscle model

We adapted an ‘Ekeberg-type’ muscle model [63] to generate torques on the joints. This model simulates muscles as a torsional spring and damper system, allowing torque control of any joint as a linear function of motor neuron (CPG output) activities driving antagonist flexor (*M*_*F*_) and extensor (*M*_*E*_) muscles controlling that joint. The torque exerted on a joint is given by the equation:

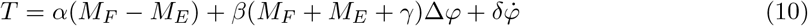

where *α, β, γ*, and *δ* represent the gain, stiffness gain, tonic stiffness, and damping coefficient, respectively [9]. Δ*ϕ* is the difference between the current angle of the joint and its resting pose. *ϕ*υ is the angular velocity of the joint. This muscle model makes it possible to control the static torque and stiffness of the joints based on optimized muscle coefficients−*α, β, γ, δ*, and Δ*ϕ*.

#### 4.3.3 CPG network and muscle parameter optimization

To identify neuromuscular network parameters that could coordinate fast and statically stable locomotion, we optimized the phase differences for each network connection, the intrinsic frequency of the oscillators, and five parameters controlling the gains and resting positions of each spring and damper muscle (i.e., *α, β, γ, δ*, and Δ*ϕ*). To simplify the problem for the optimizer, we (i) fixed ThC flexor-extensor phase differences to 180^*°*^, making them perfectly antagonistic, (ii) mirrored the phase differences from the right leg oscillators to the left leg oscillators, (iii) mirrored muscle parameters from the right joints to the left joints, and (iv) mirrored phase differences from ThC-ThC flexors to ThC-ThC extensors. Thus, a total of 63 open parameters were set by optimization: five phases between ThC CPGs (Figure 6, *A*), 12 phases between intraleg CPGs (ThC-FTi extensor/flexor, FTi-TiTa extensor/flexor per leg), 45 muscle parameters (five per joint), and one parameter (*υ*) controlling the intrinsic frequency of the oscillators. We empirically set the lower and upper bounds for the parameters so leg movements would stay stable along the boundaries (Table 6). Upper and lower bounds for the resting positions of the joints used in the muscle model were set as the first and third quartiles of measured locomotor angles. Finally, we optimized the intrinsic frequency of CPGs, denoted by *ν* in Eq. 7 to be between 6 and 10 Hz for the reasons described above.

**Table 6:**
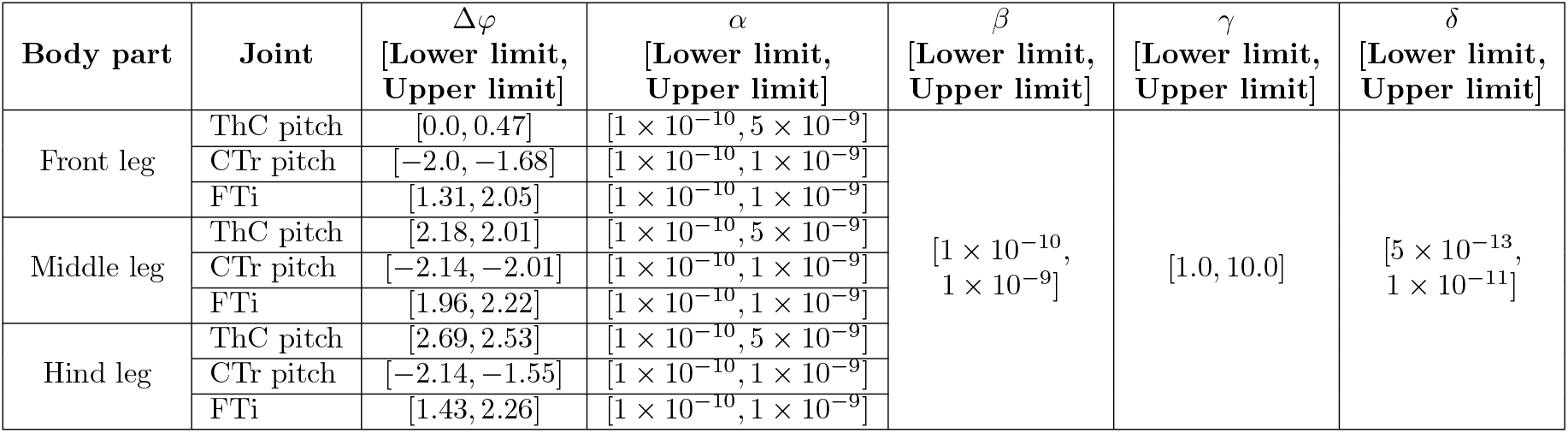
Lower and upper limits for the muscle parameters during optimization.

For parameter optimization, we used NSGA-II [67], a multi-objective genetic algorithm implemented in Python using the jMetalPy library [86]. We defined two objective functions. First, we aimed to maximize locomotor speed, as quantified by the number of spherical treadmill rotations (Equation 11) along the *Y* axis within a specific period of time. Second, we maximized static stability. In small animals like *Drosophila*, static stability is a better approximation for overall stability than dynamic stability [83]. We measured static stability by first identifying a convex hull formed by the legs in stance phase. If there were less than three legs in stance and a convex hull could not be formed, the algorithm returned −1, indicating static instability. Then, we measured the closest distance between the fly’s center of mass−dynamically calculated based on the fly’s moving body parts−and the edges of the convex hull. Finally, we obtained the minimum of all measured distances at that time step. If the center of mass was outside the convex hull, we reversed the sign of the minimum distance to indicate instability. Because the optimizer works by minimizing objective functions, we inverted the sign of speed and stability values: the most negative values meant the fastest and most stable solutions, respectively.

Four penalties were added to the objective functions. First, to make sure the model was always moving, we set a moving lower and upper threshold for the angular rotation of the ball, increasing from −0.2 *rad* to 1.0 *rad* and from 0 to 7.2 *rad* in one second, respectively. These values were determined such that the lower moving boundary was slower than the slowest reported walking speed of *Drosophila* (10 *mm/s* = 2 *rad* when the ball radius *r* is 5 *mm*) [65] and the upper moving boundary would exceed the highest reported walking speed (34 mm/s = 6.8 *rad*) [28]. Second, to avoid high torque and velocities at each joint, we set joint angular velocities to have an upper limit of 250 *rad/s*, a value measured from real fly experiments. Third, because we do not introduce physical joint limits in the model, we emulated these joint limits by setting a penalty on the difference between the joint angle range observed during kinematic replay of walking and the joint angles of individual solutions. We used this penalty to prevent joint angles from generating unrealistic movements (e.g., one full rotation around a DoF). Fourth, because the optimizer can exploit the objective function by simply leaving all legs on the ground−the highest possible stability−or can rotate the ball by using as few as two legs while the remaining legs are constantly on the ground, we introduced a penalty on duty factors. Specifically, we computed the ratio of stance phase duration to the entire epoch and penalized solutions whose duty factors for each leg were outside of the range [0.4, 0.9], based on [28].

The optimization was formulated as follows

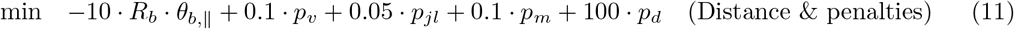

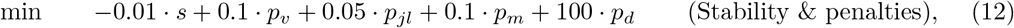

with the following penalty terms

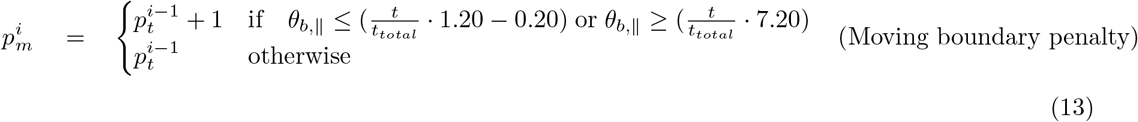

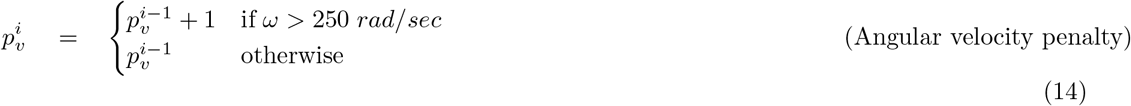

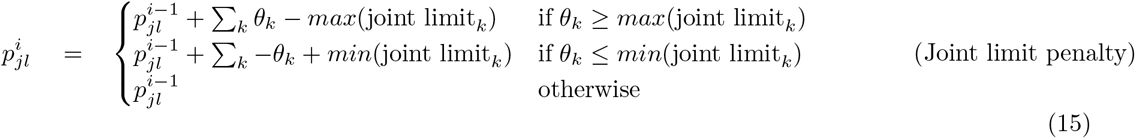

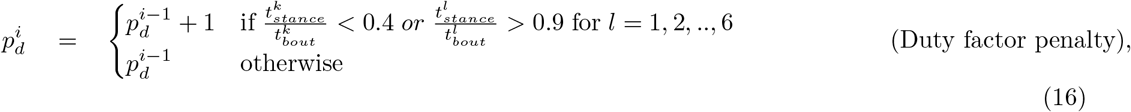

where *R*_*b*_ is the ball radius (5 mm), *θ*_*b,I*_ is the angle of the ball in the direction of walking, *t*_*tot*_ is the maximum simulation duration, *θ*_*k*_ is the angular position of the joint *k*, 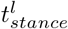 and 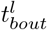 are the total times spent in stance and the entire walking epoch duration of the leg *l*. Every penalty was multiplied by its corresponding weight and added to the objective function. Objective functions were evaluated for 2 s (*t*_*total*_), a period that was sufficiently long for the model to generate locomotion. We ran 60 generations with the weights given in Equation 11 and Equation 12.

To avoid a high computational cost during optimization, we reduced the model’s complexity by removing collision shapes, like the wings and head, that were not required for locomotion, and converting joints that are not used in the simulation (see Table 4) from revolute to fixed. This model was saved as a new SDF file. Thus, we could reduce computational time and memory needed to check for collisions on unused body segments, and for the position controller to set unused joints to fixed positions. This simplification increased the speed of the simulation, allowing us to reduce the time step to 0.1 ms and to run optimization with larger populations. In the simulation, we used a spherical treadmill with a mass, radius, and friction coefficient of 54.6 *mg*, 5 *mm*, and 1.3, respectively. We additionally increased the friction coefficient of the leg segments from the default value of 0.5 to 1.0.

Each optimization generation had a population of 200 individuals. Optimization runs lasted for 60 generations, a computing time of approximately 20 hours per run on an Intel(R) Core(TM) i9-9900K CPU at 3.60GHz. Mutations occurred with a probability of 1.0 divided by number of parameters (63), and a distribution index of 20. We set the cross-over probability to 0.9 and the distribution index to 15 (for more details see [86]).

#### 4.3.4 Analysis of optimization results

After optimization, we selected three individual solutions from the last generation for deeper analysis. First, the objective functions were normalized with respect to their maximum and minimum values. Note that the signs of the objective functions were inverted. Then, solutions were selected as follows:

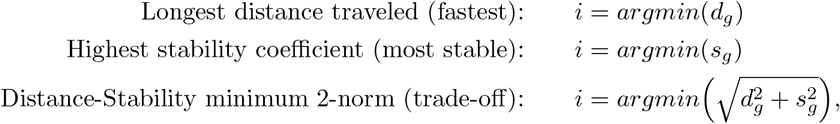

where *d*_*g*_ and *s*_*s*_ are the vectors containing the distance and stability values, respectively, from all individuals in a given generation *g*.

We plotted CPG activity patterns (as represented by the couple oscillators’ outputs), joint torques, joint angles, GRFs, and ball rotations from this final generation of solutions. GRFs were used to generate gait diagrams as previously described. Ball rotations were used to reconstruct the models’ walking paths. The distances travelled along the longitudinal (*x*) and transverse (*y*) axes were calculated from the angular displacement of the ball according to the following formula:

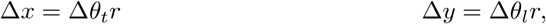

where Δ*θ*_*t*_ and Δ*θ*_*l*_ denote the angular displacement around the transverse and longitudinal axes, respectively, and *r* is the radius of the ball.

## 5 Supplementary Tables

## 6 Supplementary Figures

**Figure S1:**
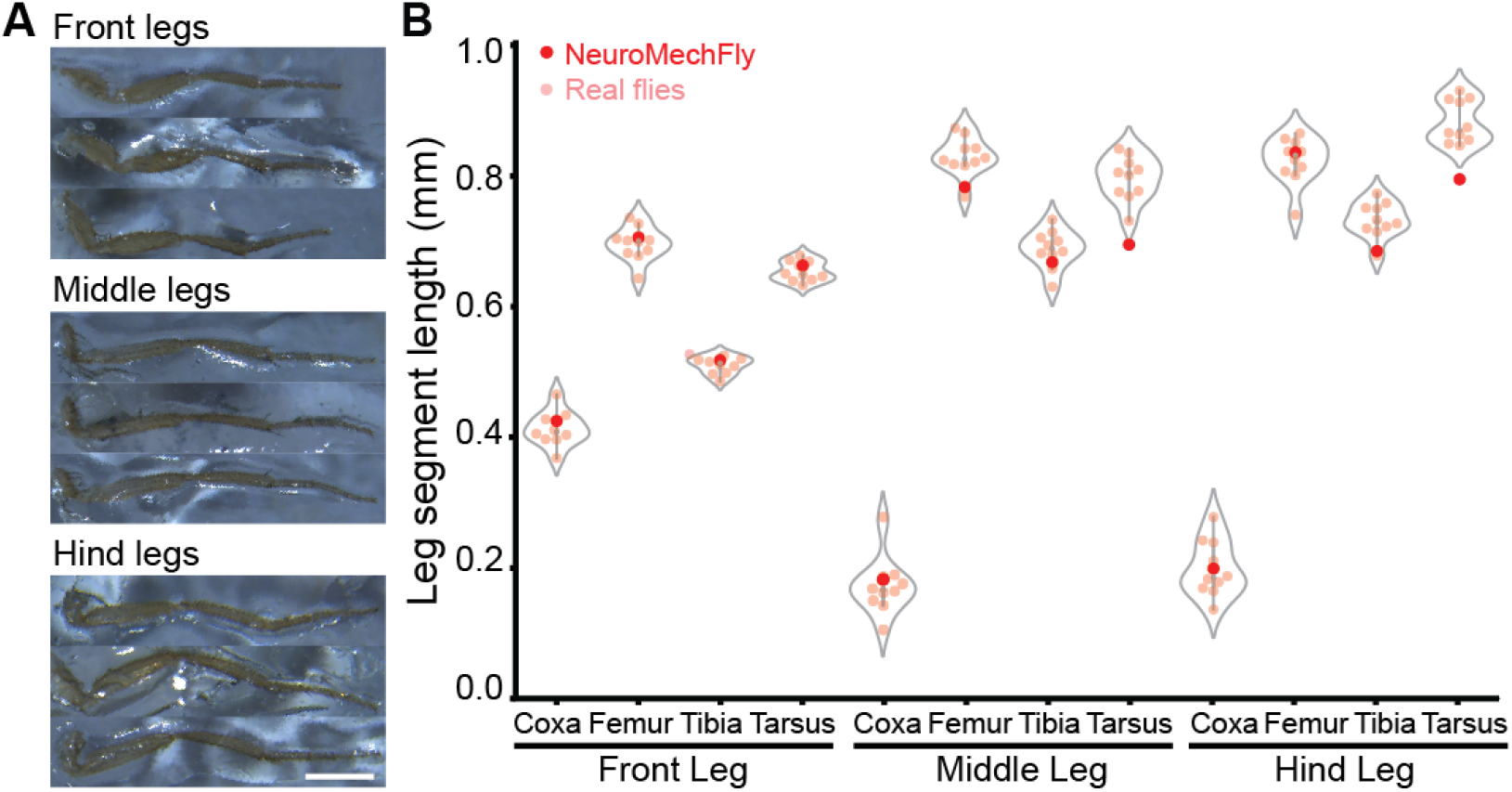
Leg segment lengths for real female *Drosophila melanogaster* and NeuroMechFly. **(A)** Legs were dissected, straightened, and fixed onto a glass slide for measurements. Scale bar is 0.5mm. **(B)** The lengths of leg segments from 1-3 dpe animals (pink) and NeuroMechFly (red) are shown. Violin plots indicate median, upper, and lower quartiles.

**Figure S2:**
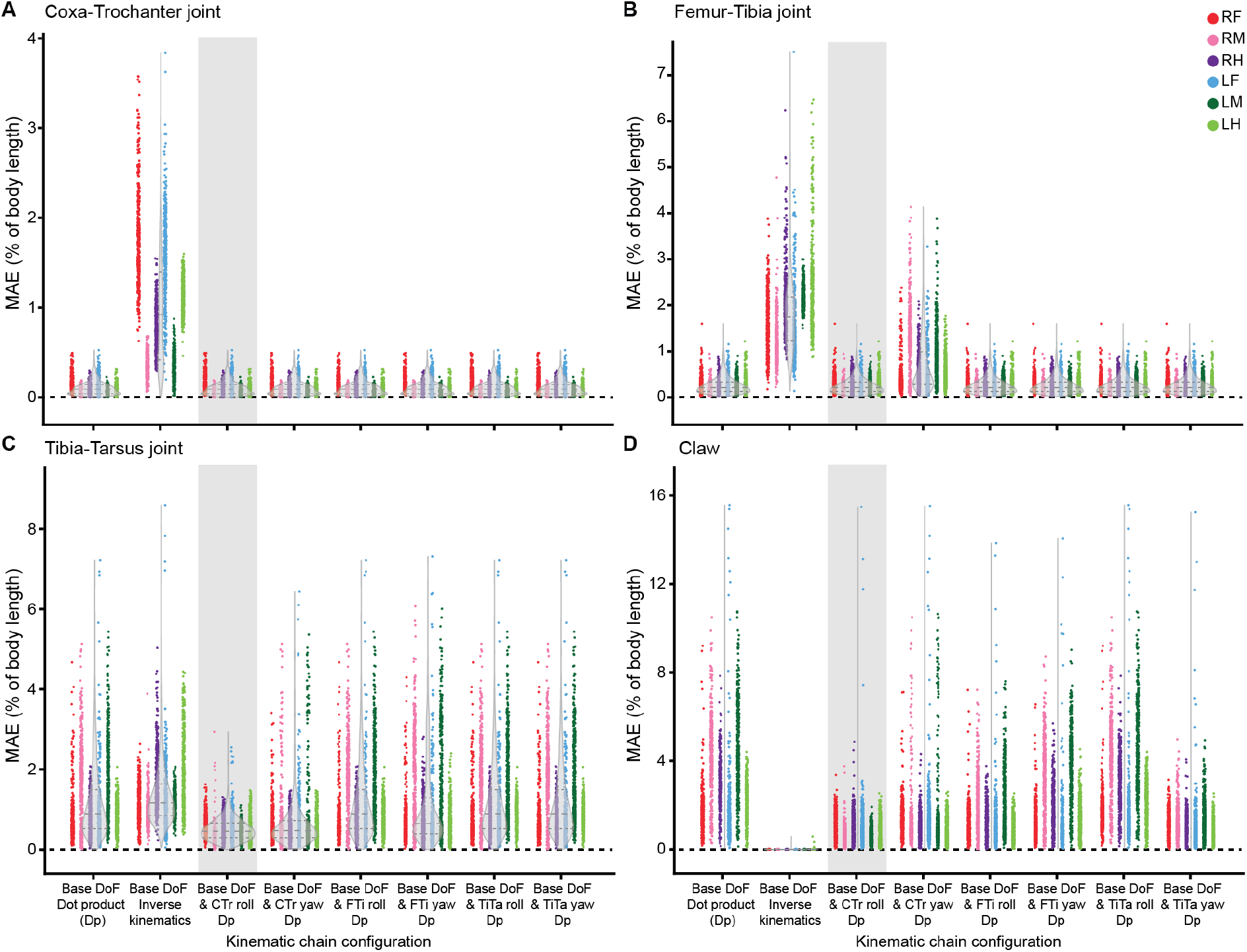
The position error for every joint in the distal leg during walking as a function of kinematic chain configuration. Body-length normalized mean absolute errors (MAE) comparing measured 3D poses and angle-derived joint positions during walking. Errors are compared among different DoF configurations for **(A)** Coxa-Trochanter joints, **(B)** Femur-Tibia joints, **(C)** Tibia-Tarsus joints, and **(D)** Claw positions. For each condition, n = 2400 samples were computed across all six legs from 4s of 100 Hz video data. Data for each leg are color-coded. ‘R’ and ‘L’ indicate right and left legs, respectively. ‘F’, ‘M’, and ‘H’ indicate front, middle, and hind legs, respectively. Violin plots indicate median, upper, and lower quartiles (dashed lines). Results from adding a coxa-trochanter roll DoF to based DoFs are highlighted in light gray.

**Figure S3:**
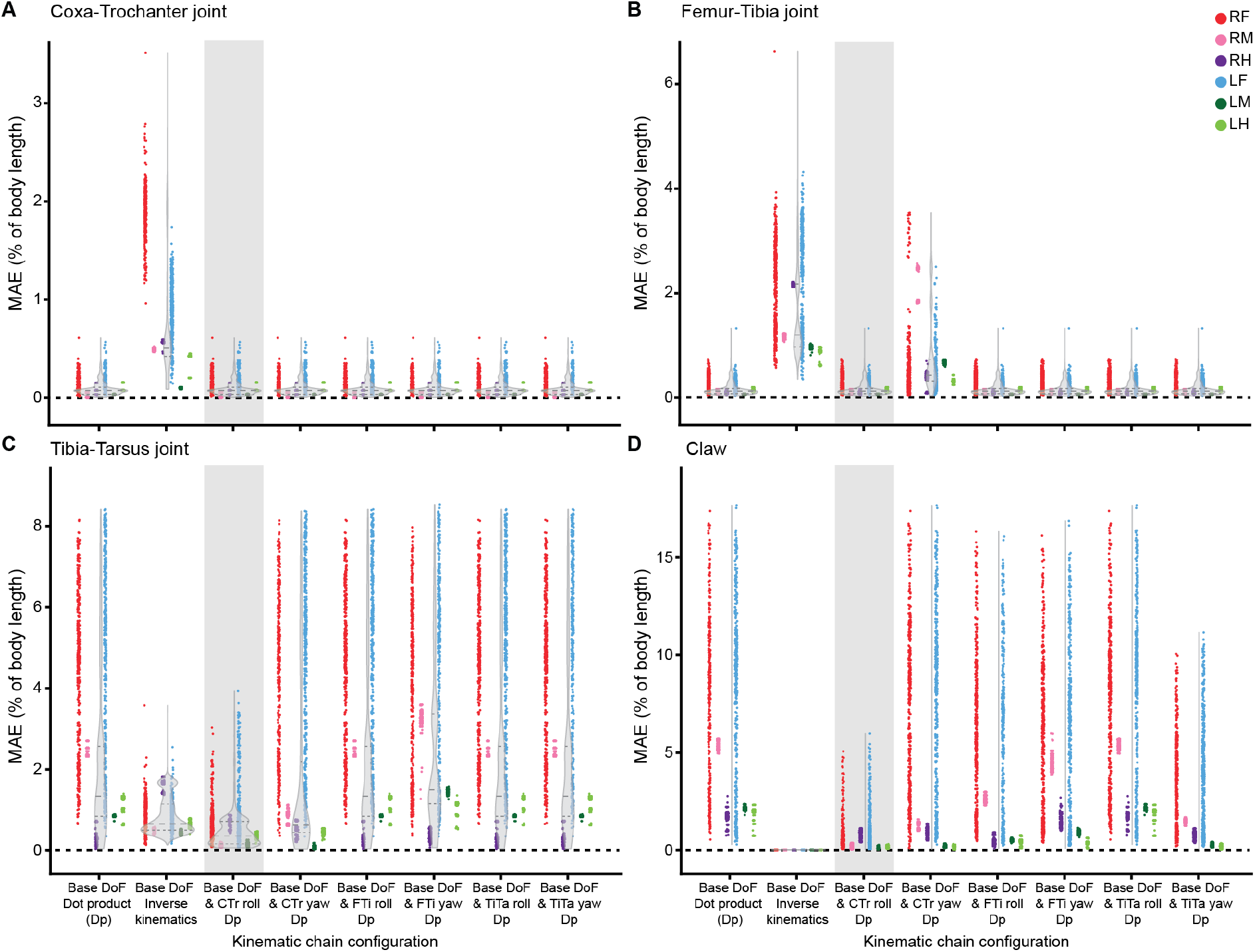
The position error for every joint in the distal leg during grooming as a function of kinematic chain configuration. Body-length normalized mean absolute errors (MAE) comparing measured 3D poses and angle-derived joint positions during grooming. Errors are compared among different DoF configurations for **(A)** Coxa-Trochanter joints, **(B)** Femur-Tibia joints, **(C)** Tibia-Tarsus joints, and **(D)** Claw positions. For each condition, n = 2400 samples were computed across all six legs from 4s of 100 Hz video data. Data for each leg are color-coded. ‘R’ and ‘L’ indicate right and left legs, respectively. ‘F’, ‘M’, and ‘H’ indicate front, middle, and hind legs, respectively. Violin plots indicate median, upper, and lower quartiles (dashed lines). Results from adding a coxa-trochanter roll DoF to based DoFs are highlighted in light gray.

**Figure S4:**
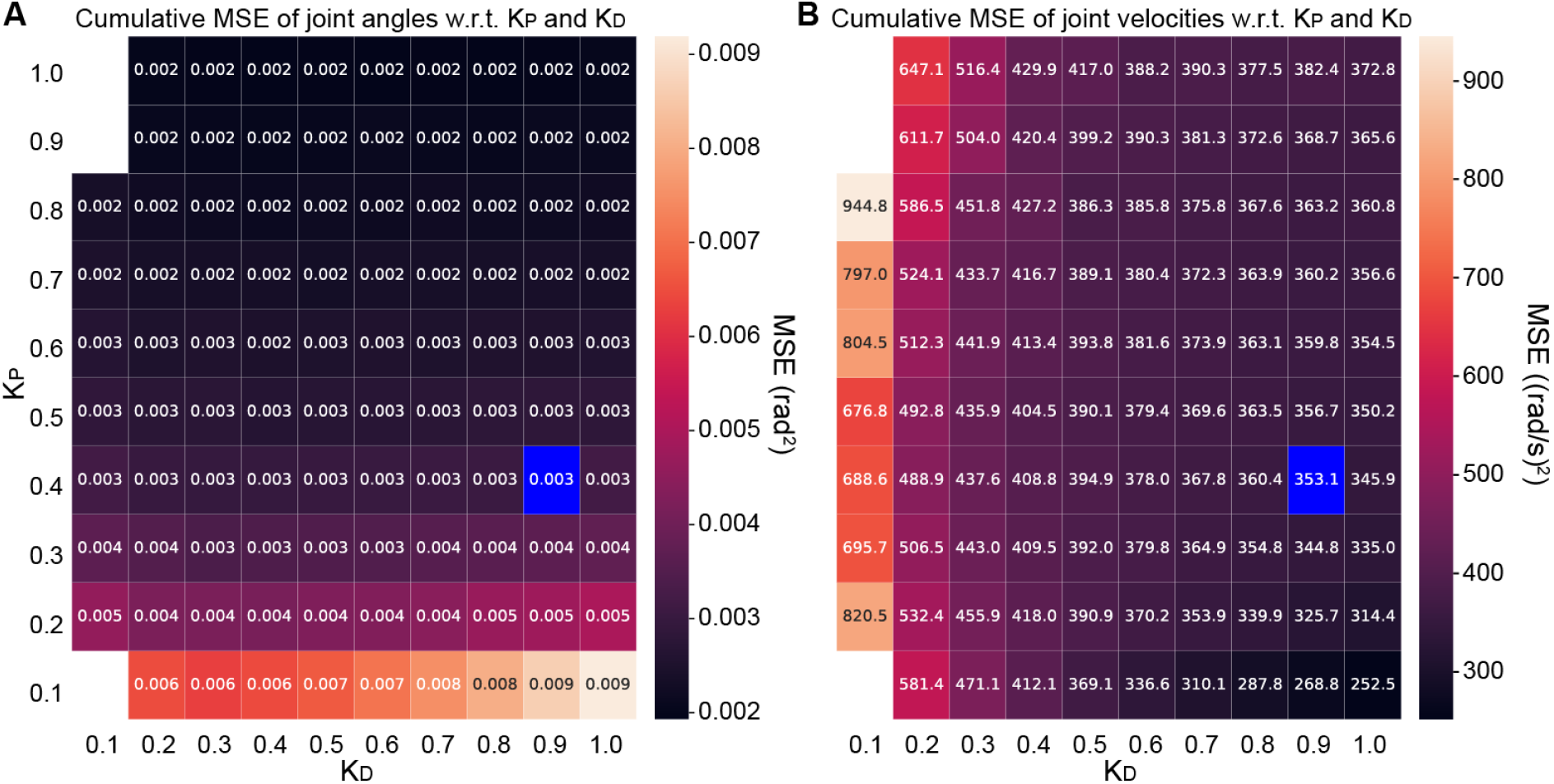
Mean squared error between tracked and simulated joint positions and velocities as a function of position and velocity gain values. MSE of **(A)** joint angles and **(B)** joint velocities as a function of derivative (*K*_*d*_) and positional gain (*K*_*p*_). Selected *K*_*p*_ and *K*_*d*_ values are indicated in blue. White areas indicate *K*_*p*_ and *K*_*d*_ pairs rendering the simulation nonfunctional.

**Figure S5:**
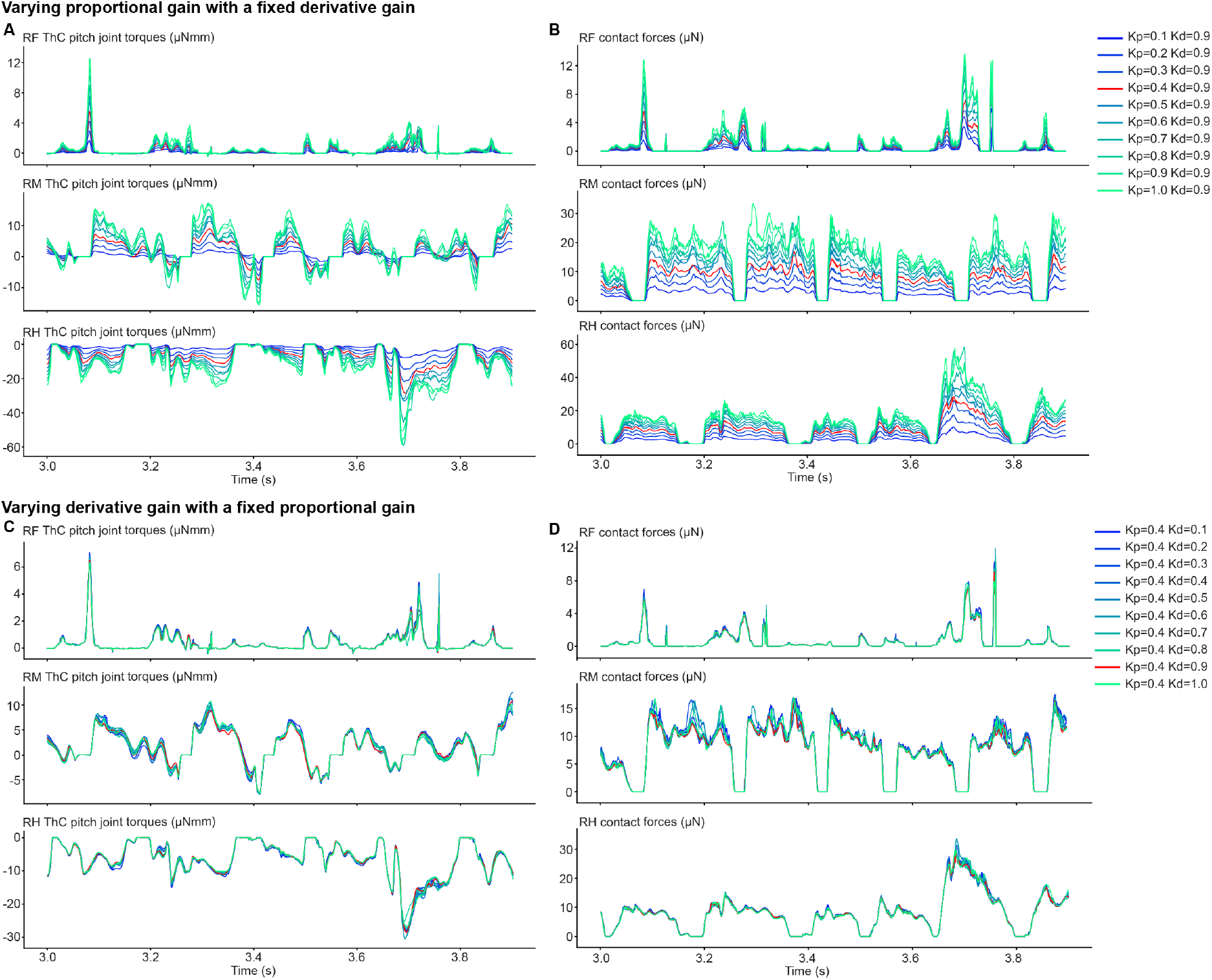
Sensitivity of estimated joint torques and contact forces to proportional and derivative gains. **(A)** Estimated torques during forward walking as a function of proportional gain (*K*_*p*_). The derivative gain (*K*_*d*_) is fixed at 0.9. Shown are measurements of ThC pitch torques for the right legs. Measurements for the contralateral legs were nearly symmetrically identical and are not shown. **(B)** Contact force measurements of the right legs during forward walking as a function of *K*_*p*_ values. Results from the selected *K*_*p*_ and *K*_*d*_ values are shown in red. **(C)** Estimated torques during forward walking as a function of derivative gain (*K*_*d*_). The proportional gain (*K*_*p*_) is fixed at 0.4. Shown are measurements of ThC pitch torques for the right legs. Measurements for the contralateral legs were nearly symmetrically identical and are not shown. **(D)** Contact force measurements of the right legs during forward walking as a function of *K*_*d*_. Results from the selected *K*_*p*_ and *K*_*d*_ values are shown in red.

**Figure S6:**
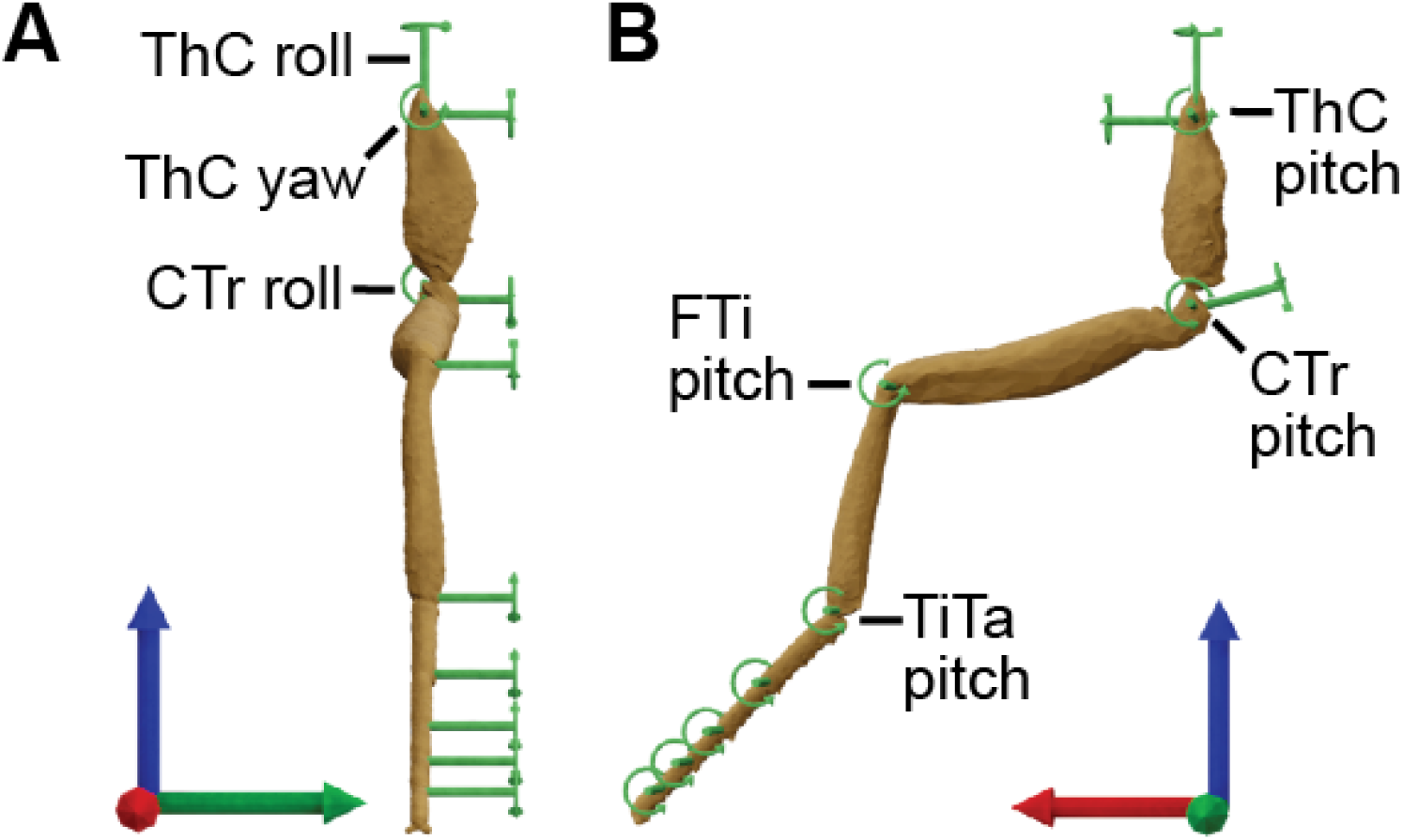
Leg joint degrees-of-freedom and their rotational axes. Each leg is composed of 11 hinge joints. Joints with more than one DoF were modeled as a union of multiple hinge joints. The left foreleg observed from **(A)** front and **(B)** side views. The global coordinate system’s x, y, and z axes are red, green, and blue, respectively.

**Figure S7:**
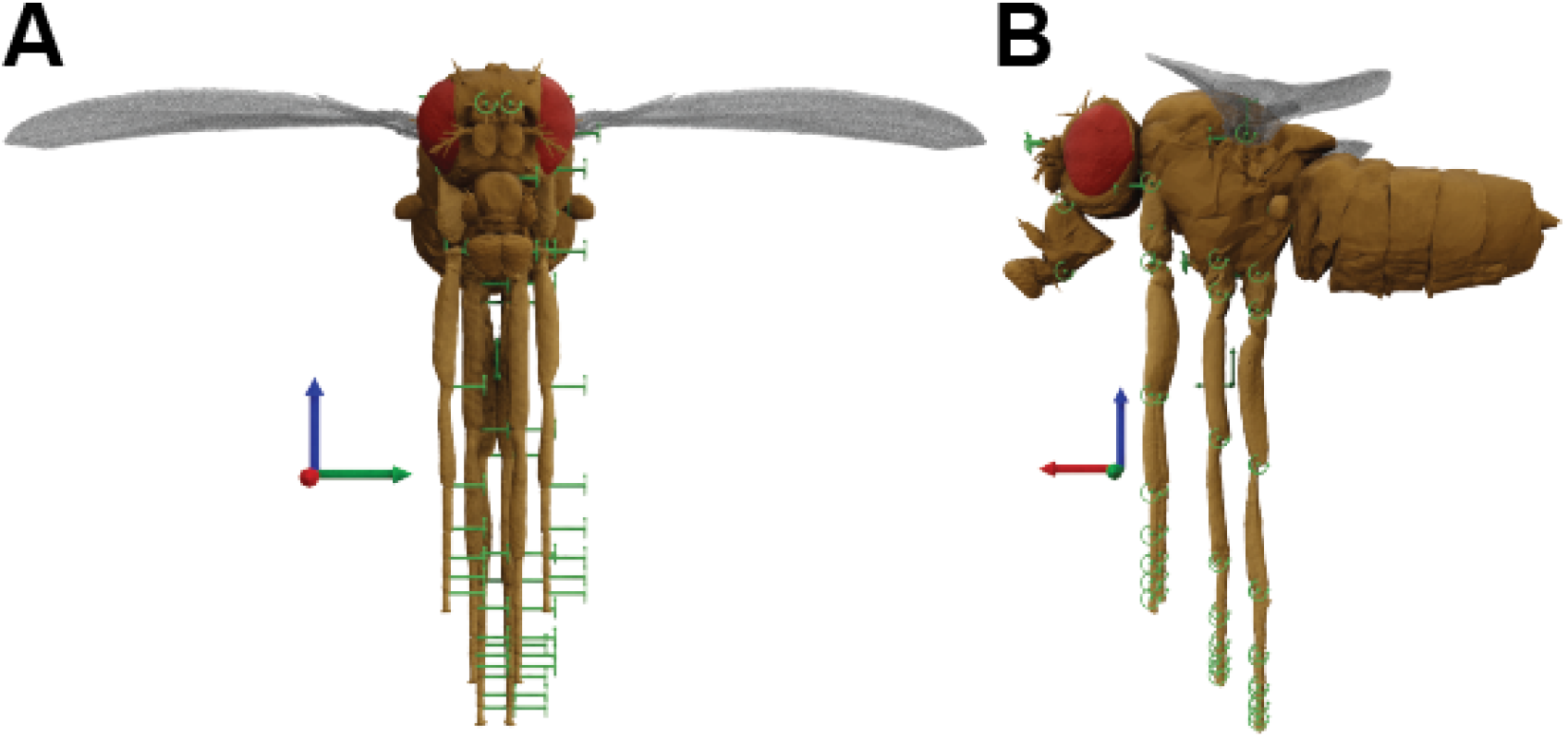
The ‘zero pose’ of NeuroMechFly. Each body segment (Table 1) is aggregated using hinge joints. Rotational axes of joints are shown. **(A)** Zero pose from **(A)** front and **(B)** side views. The global coordinate system’s x, y, and z axes are shown (red, green, and blue, respectively).

**Figure S8:**
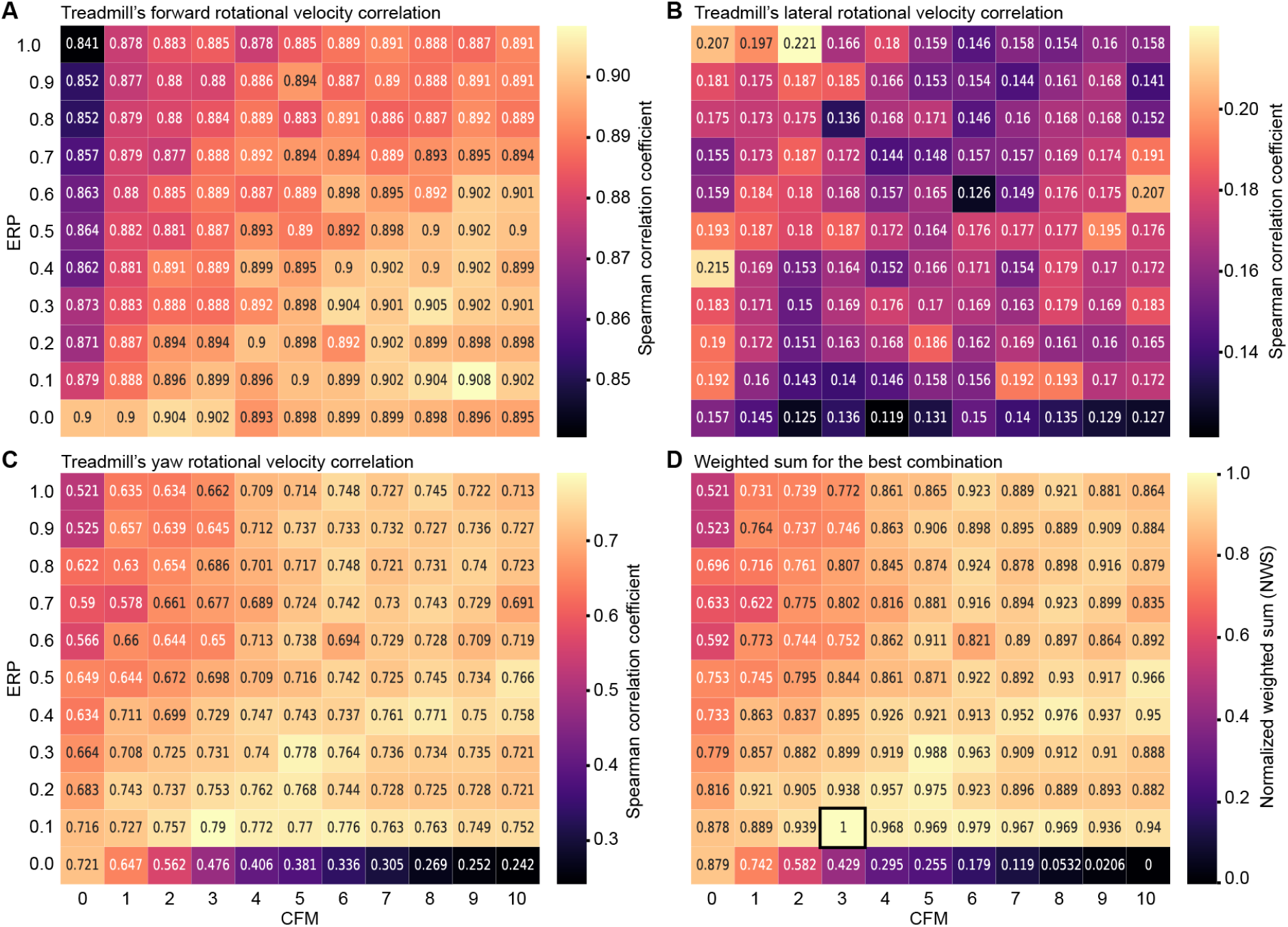
Sensitivity of simulated spherical treadmill rotation prediction accuracy during tethered walking to ERP and CFM constraint parameters. Spherical treadmill rotational velocities resulting from Kinematic Replay of walking depend on simulation constraint parameters. Shown are Spearman correlation coefficients computed between measured and estimated treadmill rotational velocities for **(A)** forward, **(B)** lateral, and **(C)** yaw axes when varying the simulation’s error reduction parameter (ERP), and the constraint force mixing (CFM). **(D)** The best combination of ERP and CFM−0.1 and 3, respectively (black outline)−was selected through a normalized weighted sum (NWS) of the correlation coefficients for each axis.

**Figure S9:**
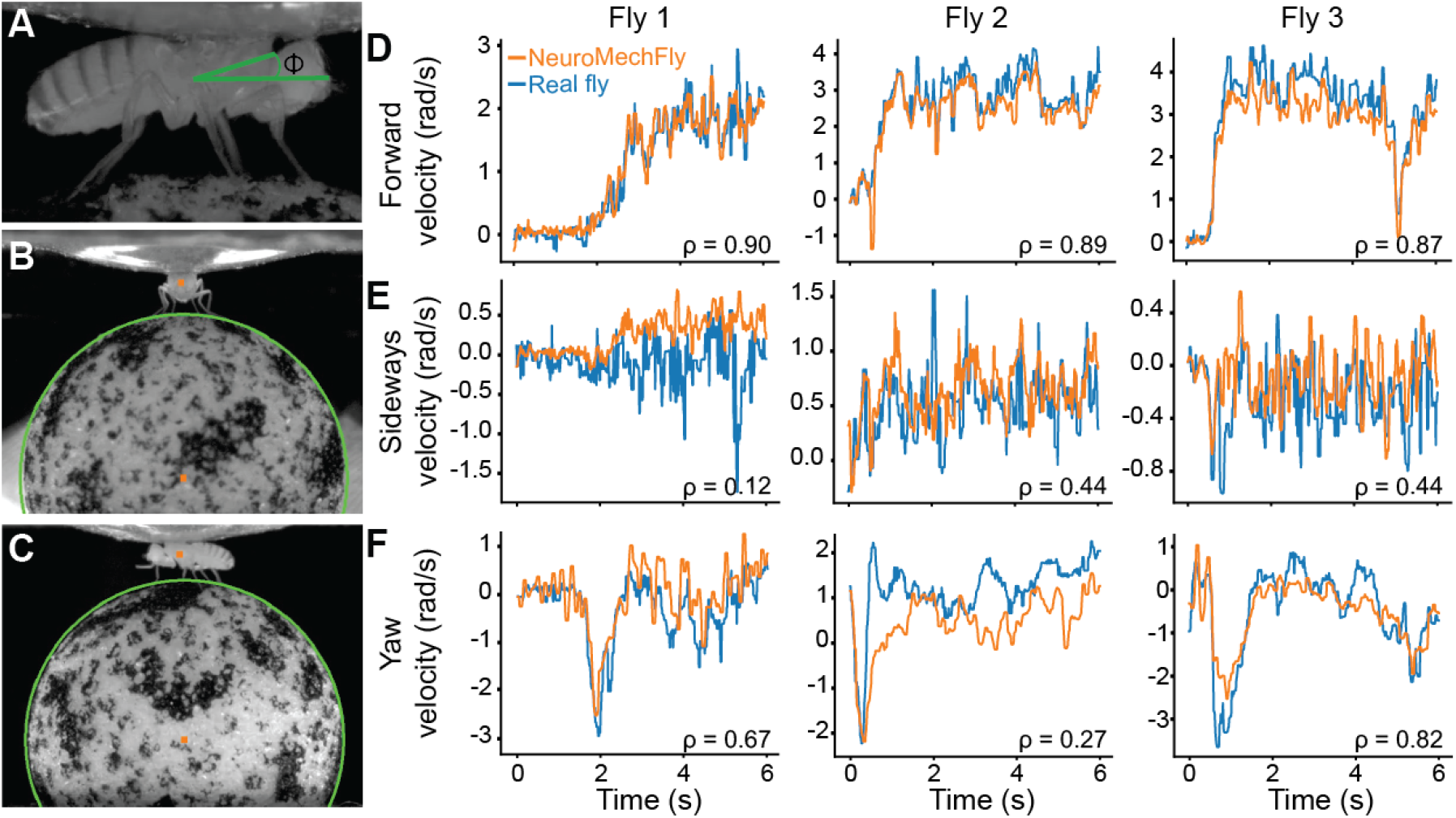
Comparing real to simulated spherical treadmill rotational velocities during tethered walking. Spherical treadmill rotations depend on a tethered fly’s **(A)** inclination (ϕ, green), **(B)** lateral, and **(C)** longitudinal positions with respect to the ball (green outlines). These positions (orange dots) were automatically detected and recreated in the simulation. Rotational velocities of the spherical treadmill generated by three real flies (blue) were compared with those generated by NeuroMechFly (orange) for **(D)** forward, **(E)** lateral, and **(F)** yaw axes. Spearman correlation coefficients (*ρ*) comparing blue and orange traces are indicated.

**Figure S10:**
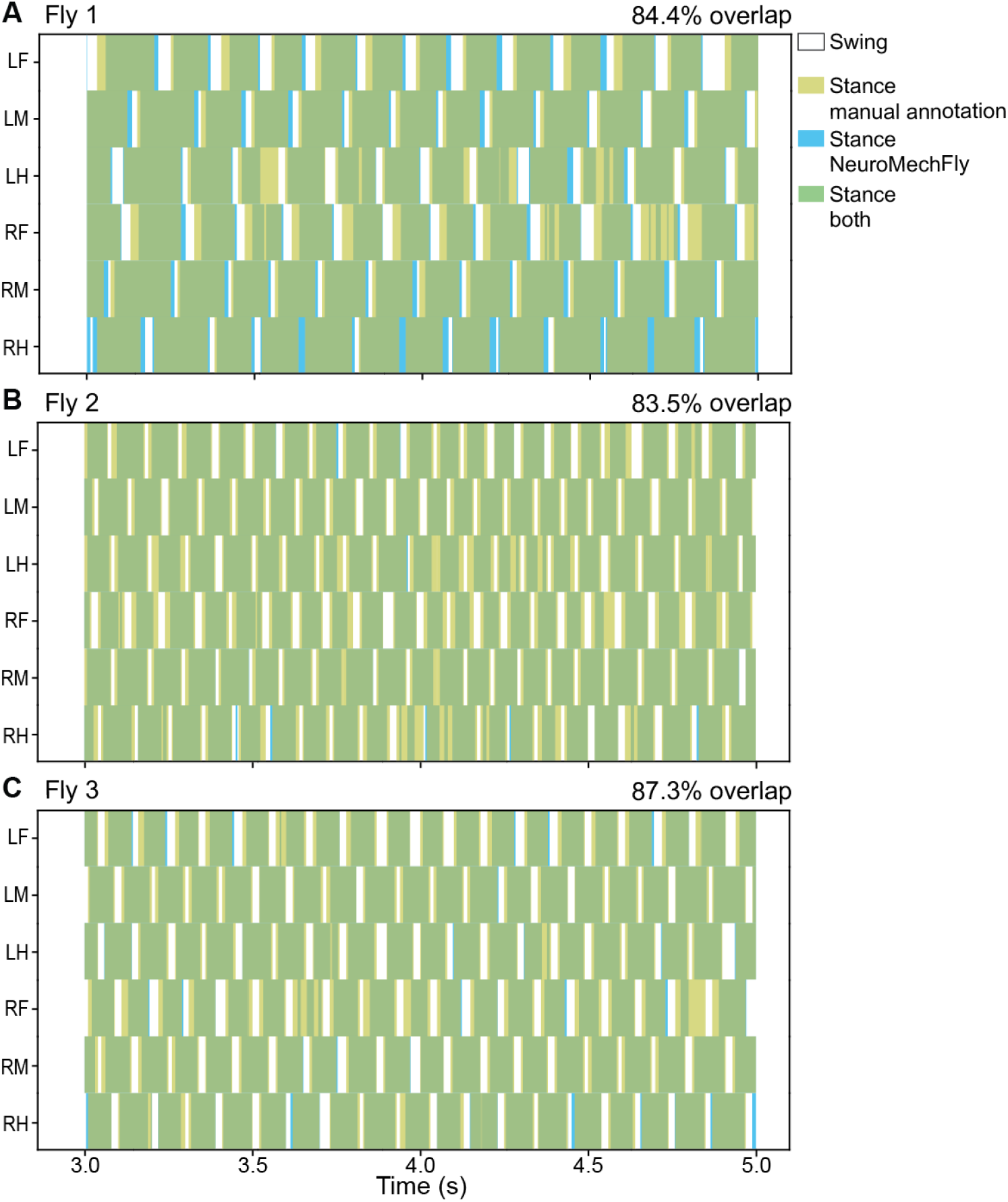
Comparing real and simulation predictions for gait diagrams during tethered walking. Gait diagrams showing manually-annotated stance phases for three real flies (**A-C**, gold) as well as those obtained from estimated ground reaction forces in NeuroMechFly (blue). Percentage of overlap in real and simulated stance phases (green) is quantified. ‘R’ and ‘L’ indicate right and left legs, respectively. ‘F’, ‘M’, and ‘H’ indicate front, middle, and hind legs, respectively.

**Figure S11:**
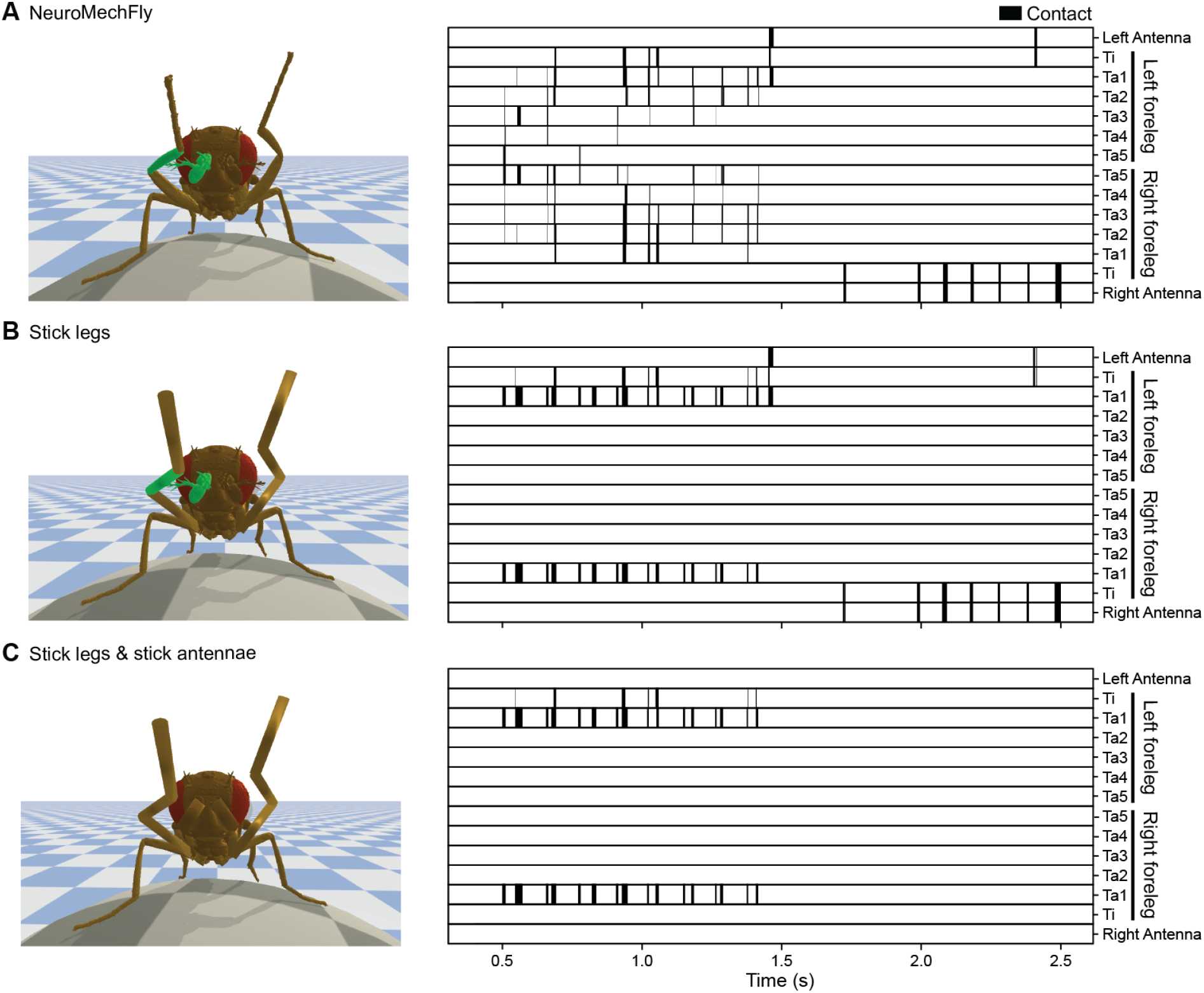
The impact of the morphological realism on estimates of leg-leg and leg-antenna contact during grooming. Collision diagrams from kinematic replay of foreleg/antennal grooming when using either **(A)** NeuroMechFly’s morphologically detailed legs and antennae, or after replacing its **(B)** forelegs, or **(C)** forelegs and antennae with simple cylinders, as in a conventional stick skeletal model.

**Figure S12:**
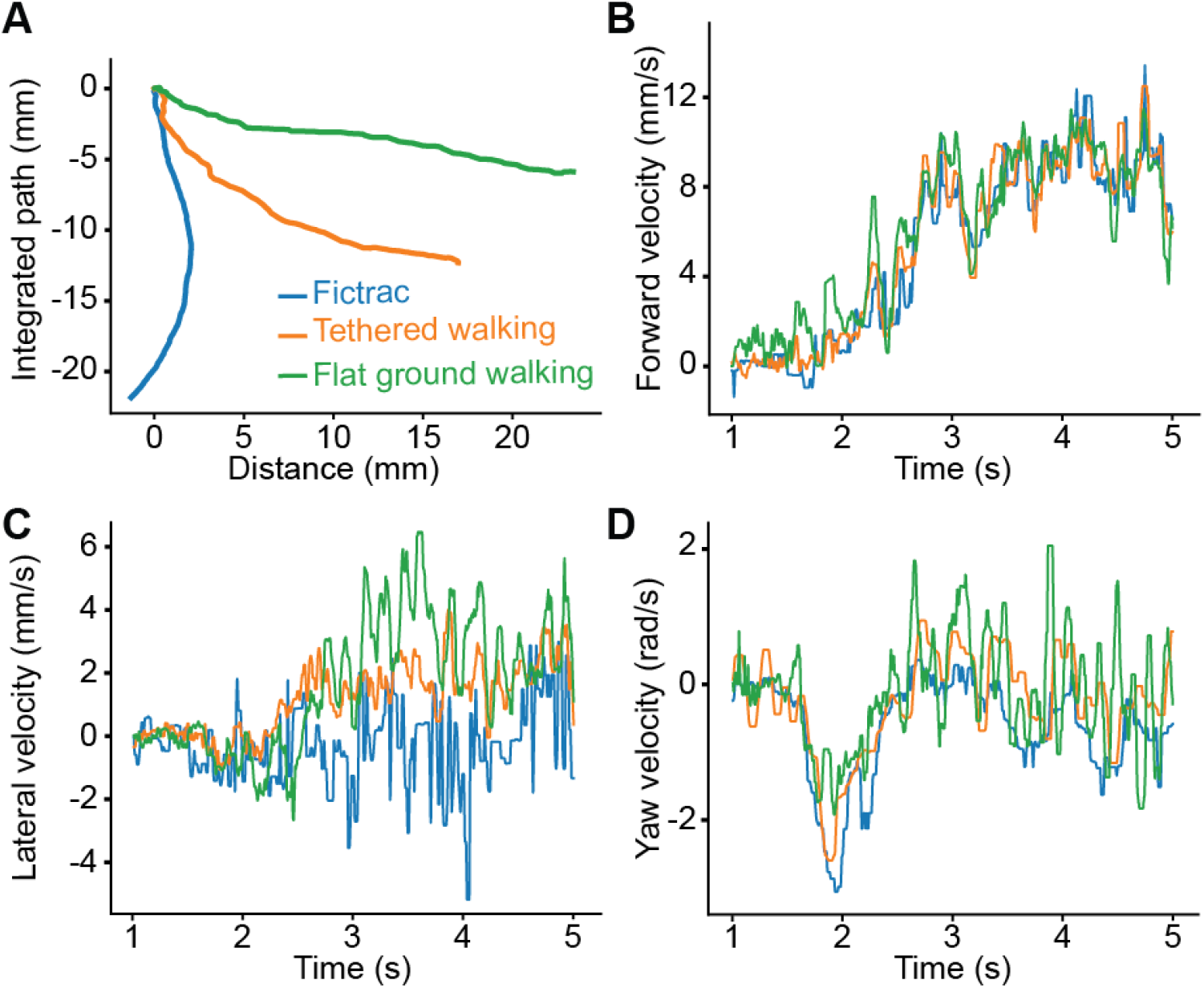
Comparison of walking paths and velocities for real tethered walking versus kinematic replay in a tethered or untethered model. Leg kinematics from a tethered walking experiment (blue) were used for kinematic replay in NeuroMechFly either tethered on a simulated spherical treadmill (orange) or freely walking on flat ground (green). Shown are resulting **(A)** integrated walking paths, as well as associated **(B)** forward, **(C)** lateral, and **(D)** yaw velocities.

**Figure S13:**
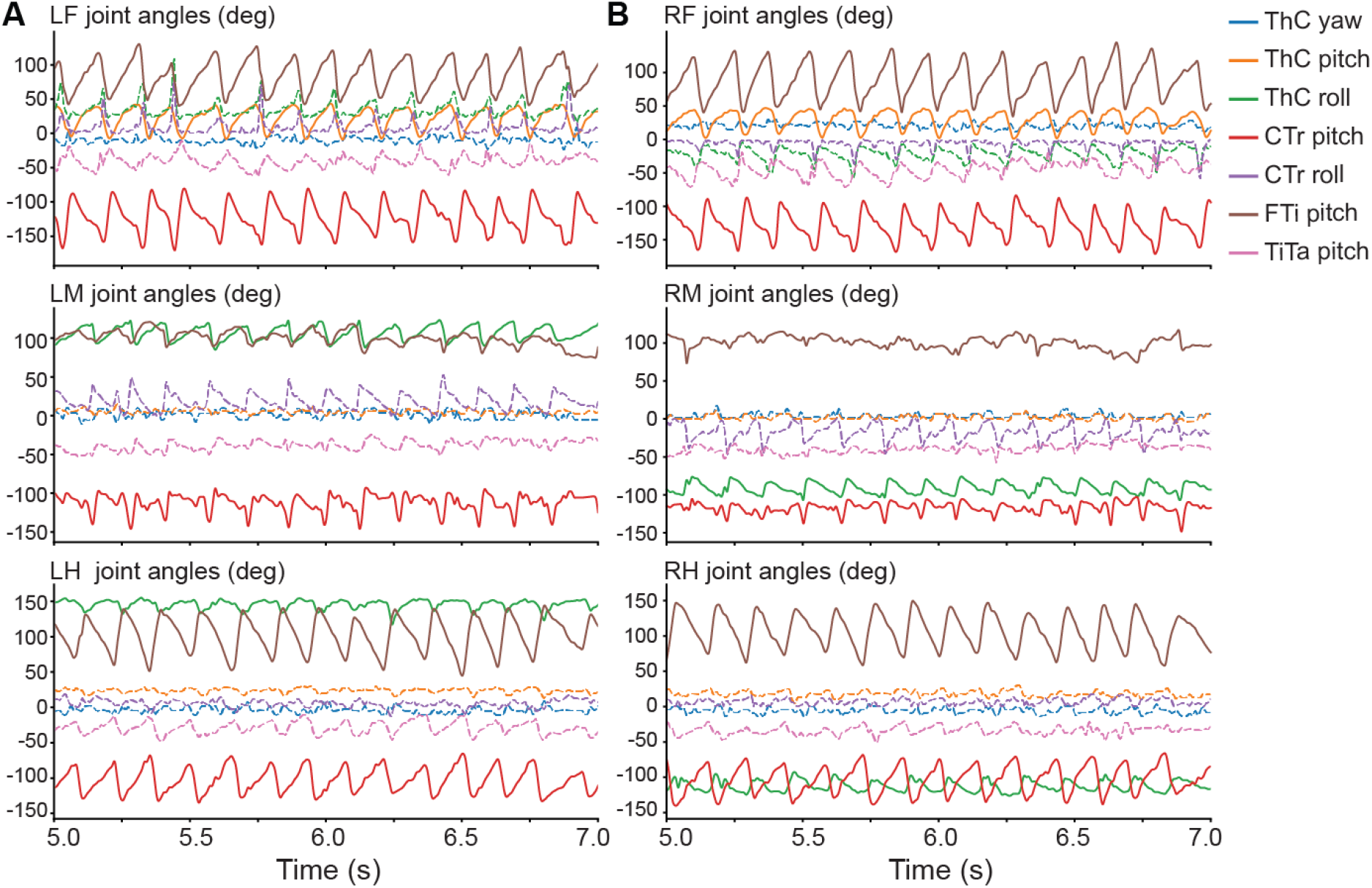
Measured joint angles during real forward walking. Joint angles for the **(A)** left and **(B)** right legs measured from a real fly during forward walking. Only the three DoFs with the highest amplitudes (solid lines) were controlled during optimization. These were: for the front legs: ThC pitch, CTr pitch, and FTi pitch; for the middle and hind legs: ThC roll, CTr pitch, and FTi pitch DoFs. The remaining four DoFs (dashed lines) for each leg did not exhibit pronounced angular changes and were fixed to their mean values during optimization.

**Figure S14:**
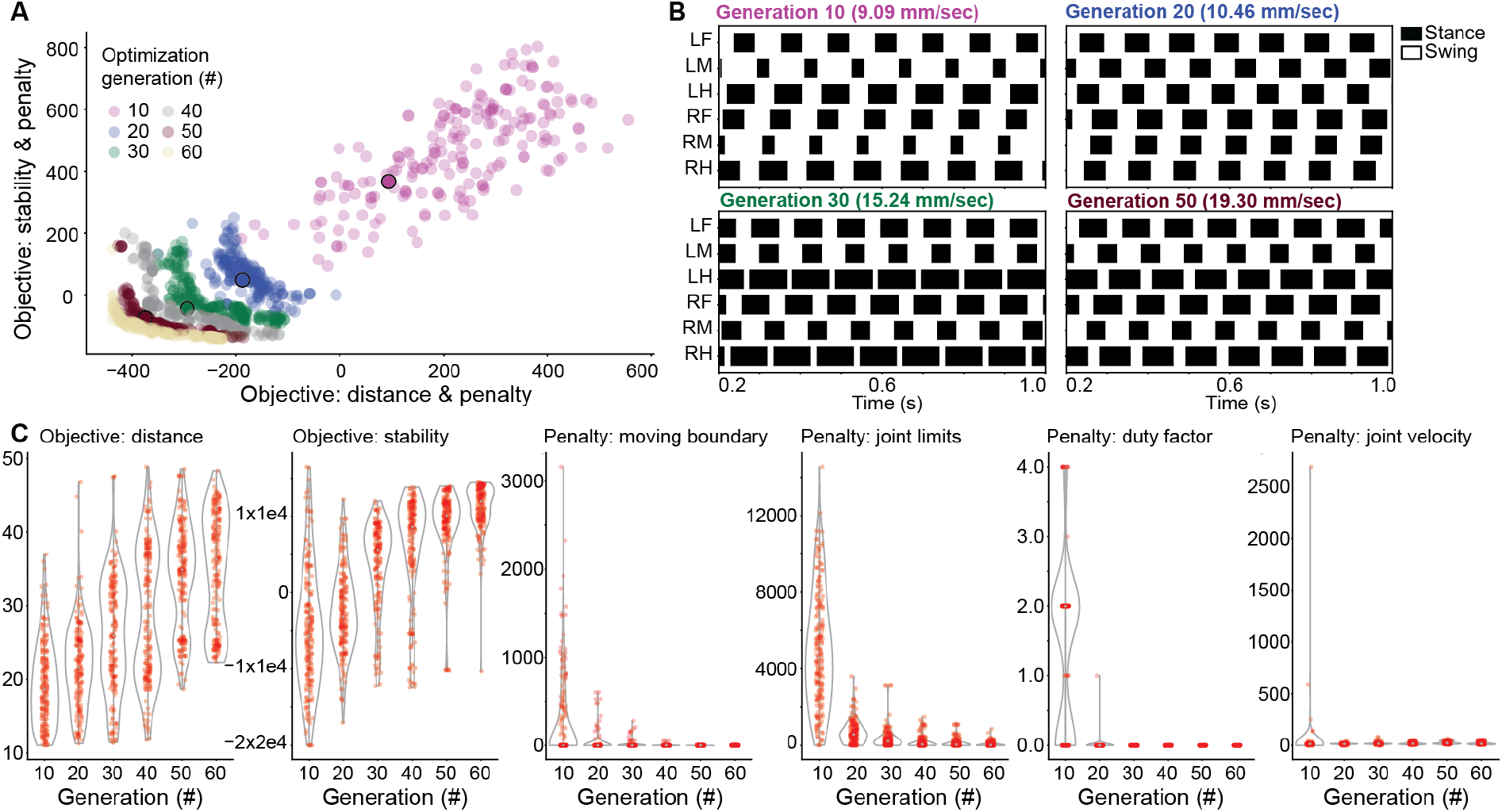
Objectives, penalties, and individual solutions over generations when optimizing for fast and statically stable tethered walking. **(A)** Pareto front approximations for six optimization generations. Later generations are more negative because the optimizer aims to minimize the distance and stability objective functions, whose signs are inverted. Four individual solutions dominated by the pareto optimal solutions were selected for more in-depth analysis (10th (purple), 20th (blue), 30th (green), and 50th (dark red); all are outlined in black). **(B)** Gait diagrams from selected solutions. Stance (black) and swing (white) phases were calculated by reading-out tarsal ground contacts for each leg. Indicated are the velocities of each solution as calculated by averaging the spherical treadmill forward velocity. **(C)** Progression of weighted objective values (shown without sign inversion) and penalties over the course of 60 generations. Objectives (distance and stability coefficients) increase across generations, while penalties decrease or converge to, or near, zero. The objective distance (mm) is the distance traveled in 2 s. The penalty duty factor is the number of legs violating the duty factor constraint. The remaining penalties are shown in Arbitrary Units.

## 7 Supplementary Videos

**Video 1: Constructing a data-driven biomechanical model of adult *Drosophila***. An adult female fly is encased in resin for x-ray microtomography. The resulting x-ray microtomography data reveals cuticle, muscles, nervous tissues, and internal organs. These data are thresholded to separate the foreground from background. Then the exoskeleton is voxelized into a 3-dimensional polygon mesh. Articulated body segments are separated from one another and then reassembled into a natural pose. Bones are added and rigged to permit actuation. Finally, textures are added to the model for visualization purposes. https://www.dropbox.com/s/pkbh4o81bdomx1x/Video1.mov?dl=0

**Video 2: Visualization of possible additional leg degrees-of-freedom**. NeuroMechFly’s left-middle leg is sequentially actuated along DoFs that are later analyzed to test their requirement for accurate replay of real fly leg kinematics. The articulated joint (e.g., ‘CTr’) and type of movement (‘roll’) are indicated. https://www.dropbox.com/s/8uhi9cyzhdntyd4/Video2.mov?dl=0

**Video 3: The effect of additional degrees-of-freedom on the accuracy of replaying forward walking**. Measured 3D poses (solid lines) and forward kinematic replay (dashed lines) for forward walking. Forward kinematics are determined either **(top-left)** using no additional degrees-of-freedom (Base DoF, dot product), **(top-middle)** instead using inverse kinematics to optimize joint angles and minimize error with only base degrees-of-freedom (Base DoF, inverse kinematics), or **(top-right and bottom row)** by adding a single new DoF (BaseDoF & ‘joint’ ‘DoF’). Legs are color-coded. https://www.dropbox.com/s/3f23rdpvz7os640/Video3.mov?dl=0

**Video 4: The effect of additional degrees-of-freedom on the accuracy of replaying foreleg/antennal grooming**. Measured 3D poses (solid lines) and forward kinematic replay (dashed lines) for foreleg/antennal grooming. Forward kinematics are determined either **(top-left)** using no additional degrees-of-freedom (Base DoF, dot product), **(top-middle)** instead using inverse kinematics to optimize joint angles and minimize error with only base degrees-of-freedom (Base DoF, inverse kinematics), or **(top-right and bottom row)** by adding a single new DoF (BaseDoF & ‘joint’ ‘DoF’). Legs are color-coded. https://www.dropbox.com/s/zv860h9ic2r8li2/Video4.mov?dl=0

**Video 5: Kinematic replay of *Drosophila* forward walking using NeuroMechFly. (top-left, ‘Raw data’)** A tethered adult fly is shown walking on a spherical treadmill. One of six synchronized camera views is shown. Data are replayed at 0.2x real time. **(bottom-left, ‘2D tracking’)** 2D poses (filled circles) and connecting ‘bones’ (lines) are superimposed for the proximal three legs. **(bottom-right, ‘3D reconstruction’)** These six 2D poses are triangulated to obtain 3D poses. Overlaid are triangulated 3D poses (solid lines) and 3D poses obtained by solving forward kinematics from joint angles (dashed lines). **(top-right, ‘Kinematic replay’)** These 3D joint angles actuate NeuroMechFly leg movements while it walks on a simulated spherical treadmill. Tarsal contacts with the ground are indicated (green). Estimated ground reaction force vectors for the proximal three legs are superimposed on the original video data **(top-left)**. https://www.dropbox.com/s/iieuwgmx8bazzmd/Video5.mov?dl=0

**Video 6: Kinematic replay of *Drosophila* foreleg/antennal grooming using NeuroMechFly. (top-left, ‘Raw data’)** A tethered adult fly is shown grooming on a spherical treadmill. One of six synchronized camera views is shown. Data are replayed at 0.2x real time. **(bottom-left, ‘2D tracking’)** 2D poses (filled circles) and connecting ‘bones’ (lines) are superimposed for the proximal three legs. **(bottom-right, ‘3D reconstruction’)** These six 2D poses are triangulated to obtain 3D poses. Overlaid are triangulated 3D poses (solid lines) and 3D poses obtained by solving forward kinematics from joint angles (dashed lines). **(top-right, ‘Kinematic replay’)** These joint angles actuate NeuroMechFly leg movements while it grooms on a simulated spherical treadmill. Leg segments and antennal collisions are indicated (green). Estimated collision force vectors for the front legs and antennae are subsequently superimposed on the original video data **(top-left)**. https://www.dropbox.com/s/m3j6wfevzenhfkn/Video6.mov?dl=0

**Video 7: The influence of leg and antenna morphological detail on collision predictions. (top-left, ‘Raw data’)** Real fly grooming as recorded from the front camera. **(top-right, ‘NeuroMechFly’)** NeuroMechFly performing kinematic replay of grooming. **(bottom-left, ‘Stick model legs’)** NeuroMechFly with stick legs but detailed antennae. **(bottom-right, ‘Stick model legs and antennae’)** NeuroMechFly with stick legs and stick antennae. https://www.dropbox.com/s/7wpnf2a8s4pzi65/Video7.mov?dl=0

**Video 8: Kinematic replay of tethered *Drosophila* forward walking using NeuroMechFly on flat terrain without body support. (Right)** Pose estimates obtained from a real tethered fly walking on a spherical treadmill are replayed in NeuroMechFly as it walks untethered on flat terrain without body support. **(Left)** Integrated paths are shown for tethered (orange) and flat ground (green) scenarios. https://www.dropbox.com/s/e7qvz4tm1exhefl/Video8.mov?dl=0

**Video 9: Forward walking across optimization generations**. Forward walking for four solutions shown across optimization generations 15, 30, 45 and 60. Tarsal contacts with the ground are indicated (green). Videos are replayed at 0.1x real time. Solutions shown are: (top-left) a random individual, (top-right) the fastest individual (i.e., with the longest distance traveled), (bottom-left) the most stable individual, and (bottom-right) the best trade-off achieving both high speed and static stability. https://www.dropbox.com/s/lizgd3ss2yftlxb/Video9.mov?dl=0

**Video 10: Replaying real tethered walking kinematics on flat terrain and applying external perturbations**. Pose estimates obtained from a real tethered fly walking on a spherical treadmill are replayed in NeuroMechFly as it walks untethered on flat terrain without body support. Simulated spheres are projected at the model to illustrate perturbations and the possibility of using more complex physical environments in PyBullet. https://www.dropbox.com/s/ae6zrejhddwduun/Video10.mov?dl=0

## 8 Code and data availability

Data are available at

https://doi.org/10.7910/DVN/Y3TAEC

Code, and documentation are available at:

https://github.com/NeLy-EPFL/NeuroMechFly https://nely-epfl.github.io/NeuroMechFly

## 10 Acknowledgments

We thank Stéphanie Clerc Rosset and Graham Knott (Biological Electron Microscopy Facility, EPFL, Lausanne, Switzerland) for preparing *Drosophila melanogaster* samples for X-ray microtomography. We thank Halla Sigurthorsdottir for early work on fly leg degrees-of-freedom.

## 11 Author Contributions

V.L.R. - Conceptualization, Methodology, Software, Validation, Formal Analysis, Investigation, Data Curation, Validation, Writing – Original Draft Preparation, Writing – Review & Editing, Visualization.

S.T.R. - Conceptualization, Methodology, Software, Validation, Writing – Review & Editing, Visualization.

P.G.O. - Conceptualization, Methodology, Software, Validation, Formal Analysis, Investigation, Data Curation, Writing – Review & Editing, Visualization.

J.A. - Conceptualization, Methodology, Software, Validation, Writing – Review & Editing.

A.J.I. - Conceptualization, Methodology, Resources, Writing - Review & Editing, Supervision, Project Administration, Funding Acquisition.

P.R. - Conceptualization, Methodology, Resources, Writing – Original Draft Preparation, Writing - Review & Editing, Supervision, Project Administration, Funding Acquisition.

## References

[1] Chiel, H. J. & Beer, R. D. The brain has a body: adaptive behavior emerges from interactions of nervous system, body and environment. Trends in neurosciences 20, 553–557 (1997).

[2] Webb, B. A framework for models of biological behaviour. International journal of neural systems 9, 375–381 (1999).

[3] Pearson, K., Ekeberg, Ö. & Büschges, A. Assessing sensory function in locomotor systems using neuro-mechanical simulations. Trends in neurosciences 29, 625–631 (2006).

[4] Prilutsky, B. I. & Edwards, D. H. Neuromechanical modeling of posture and locomotion (2015).

[5] Seth, A. et al. OpenSim: Simulating musculoskeletal dynamics and neuromuscular control to study human and animal movement. PloS computational biology 14 (2018).

[6] Einevoll, G. T. et al. The scientific case for brain simulations. Neuron 102, 735–744 (2019).

[7] Sigvardt, K. A. & Miller, W. L. Analysis and modeling of the locomotor central pattern generator as a network of coupled oscillators. Annals of the New York Academy of Sciences 860, 250–265 (1998).

[8] Lansner, A., Hellgren Kotaleski, J. & Grillner, S. Modeling of the spinal neuronal circuitry underlying locomotion in a lower vertebratea. Annals of the New York Academy of Sciences 860, 239–249 (1998).

[9] Ijspeert, A. J. A connectionist central pattern generator for the aquatic and terrestrial gaits of a simulated salamander. Biological cybernetics 84, 331–348 (2001).

[10] Rybak, I. A., Dougherty, K. J. & Shevtsova, N. A. Organization of the mammalian locomotor CPG: review of computational model and circuit architectures based on genetically identified spinal interneurons. ENeuro 2 (2015).

[11] Ekeberg, Ö., Blümel, M. & Büschges, A. Dynamic simulation of insect walking. Arthropod structure & development 33, 287–300 (2004).

[12] Toth, T. I., Schmidt, J., Büschges, A. & Daun-Gruhn, S. A neuro-mechanical model of a single leg joint highlighting the basic physiological role of fast and slow muscle fibres of an insect muscle system. PloS one 8 (2013).

[13] Toth, T. I., Grabowska, M., Schmidt, J., Büschges, A. & Daun-Gruhn, S. A neuro-mechanical model explaining the physiological role of fast and slow muscle fibres at stop and start of stepping of an insect leg. PloS one 8 (2013).

[14] Schilling, M., Hoinville, T., Schmitz, J. & Cruse, H. Walknet, a bio-inspired controller for hexapod walking. Biological cybernetics 107, 397–419 (2013).

[15] Szczecinski, N. S., Brown, A. E., Bender, J. A., Quinn, R. D. & Ritzmann, R. E. A neuromechanical simulation of insect walking and transition to turning of the cockroach Blaberus discoidalis. Biological cybernetics 108, 1–21 (2014).

[16] Proctor, J., Kukillaya, R. & Holmes, P. A phase-reduced neuro-mechanical model for insect locomotion: feed-forward stability and proprioceptive feedback. Philosophical Transactions of the Royal Society A: Mathematical, Physical and Engineering Sciences 368, 5087–5104 (2010).

[17] Szczecinski, N. S., Martin, J. P., Bertsch, D. J., Ritzmann, R. E. & Quinn, R. D. Neurome-chanical model of praying mantis explores the role of descending commands in pre-strike pivots. Bioinspiration & biomimetics 10 (2015).

[18] Guo, S., Lin, J., Wöhrl, T. & Liao, M. A Neuro-Musculo-Skeletal Model for Insects With Data-driven Optimization. Scientific reports 8, 1–11 (2018).

[19] Szigeti, B. et al. OpenWorm: an open-science approach to modeling Caenorhabditis elegans. Frontiers in computational neuroscience 8, 137 (2014).

[20] Izquierdo, E. J. & Beer, R. D. From head to tail: a neuromechanical model of forward locomotion in Caenorhabditis elegans. Philosophical Transactions of the Royal Society B: Biological Sciences 373 (2018).

[21] Loveless, J., Lagogiannis, K. & Webb, B. Modelling the mechanics of exploration in larval Drosophila. PloS computational biology 15 (2019).

[22] Merel, J. et al. Deep neuroethology of a virtual rodent. arXiv (2019).

[23] Isakov, A. et al. Recovery of locomotion after injury in Drosophila melanogaster depends on proprioception. Journal of Experimental Biology 219, 1760–1771 (2016).

[24] Ramdya, P. et al. Climbing favours the tripod gait over alternative faster insect gaits. Nature communications 8 (2017).

[25] Seeds, A. M. et al. A suppression hierarchy among competing motor programs drives sequential grooming in Drosophila. eLife 3 (2014).

[26] Pavlou, H. J. & Goodwin, S. F. Courtship behavior in Drosophila melanogaster: towards a ‘courtship connectome’. Current Opinion in Neurobiology 23, 76–83 (2013).

[27] Fry, S. N., Sayaman, R. & Dickinson, M. H. The aerodynamics of free-flight maneuvers in Drosophila. Science 300, 495–498 (2003).

[28] Mendes, C. S., Bartos, I., Akay, T., Márka, S. & Mann, R. S. Quantification of gait parameters in freely walking wild type and sensory deprived Drosophila melanogaster. eLife 2 (2013).

[29] Wosnitza, A., Bockemühl, T., Dübbert, M., Scholz, H. & Büschges, A. Inter-leg coordination in the control of walking speed in Drosophila. Journal of Experimental Biology 216, 480–491 (2013).

[30] Pick, S. & Strauss, R. Goal-driven behavioral adaptations in gap-climbing Drosophila. Current Biology 15, 1473–1478 (2005).

[31] Pereira, T. D. et al. Fast animal pose estimation using deep neural networks. Nature methods 16, 117–125 (2019).

[32] Mathis, A. et al. DeepLabCut: markerless pose estimation of user-defined body parts with deep learning. Nature neuroscience 21, 1281–1289 (2018).

[33] Günel, S. et al. DeepFly3D, a deep learning-based approach for 3D limb and appendage tracking in tethered, adult Drosophila. eLife 8 (2019).

[34] Gosztolai, A. et al. LiftPose3D, a deep learning-based approach for transforming two-dimensional to three-dimensional poses in laboratory animals. Nature methods 18, 975–981 (2021).

[35] Jenett, A. et al. A GAL4-driver line resource for Drosophila neurobiology. Cell reports 2, 991–1001 (2012).

[36] Seelig, J. D. et al. Two-photon calcium imaging from head-fixed Drosophila during optomotor walking behavior. Nature methods 7, 535 (2010).

[37] Maimon, G., Straw, A. D. & Dickinson, M. H. Active flight increases the gain of visual motion processing in Drosophila. Nature neuroscience 13, 393 (2010).

[38] Chen, C.-L. et al. Imaging neural activity in the ventral nerve cord of behaving adult Drosophila. Nature communications 9 (2018).

[39] Hermans, L. et al. Long-term imaging of the ventral nerve cord in behaving adult Drosophila. bioRxiv (2021).

[40] Phelps, J. S. et al. Reconstruction of motor control circuits in adult Drosophila using automated transmission electron microscopy. Cell 184, 759–774 (2021).

[41] Scheffer, L. K. et al. A Connectome and Analysis of the Adult Drosophila Central Brain. bioRxiv (2020).

[42] Coumans, E. Bullet physics simulation. In ACM SIGGRAPH 2015 Courses (2015).

[43] Lewiner, T., Lopes, H., Vieira, A. W. & Tavares, G. Efficient Implementation of Marching Cubes’ Cases with Topological Guarantees. Journal of Graphics Tools 8, 1–15 (2003).

[44] Soler, C., Daczewska, M., Da Ponte, J. P., Dastugue, B. & Jagla, K. Coordinated development of muscles and tendons of the Drosophila leg. Development 131, 6041–6051 (2004).

[45] Sink, H. Muscle development in drosophila (2006).

[46] Cruse, H., Dürr, V. & Schmitz, J. Insect walking is based on a decentralized architecture revealing a simple and robust controller. Philosophical Transactions of the Royal Society A: Mathematical, Physical and Engineering Sciences 365, 221–250 (2007).

[47] Loper, M., Mahmood, N., Romero, J., Pons-Moll, G. & Black, M. J. SMPL: A skinned multi-person linear model. ACM transactions on graphics 34, 1–16 (2015).

[48] Zuffi, S., Kanazawa, A., Jacobs, D. W. & Black, M. J. 3D menagerie: Modeling the 3D shape and pose of animals. In Proceedings of the IEEE conference on computer vision and pattern recognition, 6365–6373 (2017).

[49] Li, S. et al. Deformation-aware Unpaired Image Translation for Pose Estimation on Laboratory Animals. arXiv (2020).

[50] Mu, J., Qiu, W., Hager, G. D. & Yuille, A. L. Learning from Synthetic Animals. In Proceedings of the IEEE/CVF Conference on Computer Vision and Pattern Recognition, 12386–12395 (2020).

[51] Bolanõs, L. A. et al. A three-dimensional virtual mouse generates synthetic training data for behavioral analysis. Nature methods (2021).

[52] Watson, J. T., Ritzmann, R. E., Zill, S. N. & Pollack, A. J. Control of obstacle climbing in the cockroach, Blaberus discoidalis. I. Kinematics. Journal of Comparative Physiology A 188, 39–53 (2002).

[53] Frantsevich, L. & Wang, W. Gimbals in the insect leg. Arthropod structure & development 38, 16–30 (2009).

[54] Bender, J. A., Simpson, E. M. & Ritzmann, R. E. Computer-assisted 3D kinematic analysis of all leg joints in walking insects. PloS one 5 (2010).

[55] Zill, S. N. et al. Effects of force detecting sense organs on muscle synergies are correlated with their response properties. Arthropod structure & development 46, 564–578 (2017).

[56] Cofer, D., Cymbalyuk, G., Heitler, W. J. & Edwards, D. H. Neuromechanical simulation of the locust jump. Journal of Experimental Biology 213, 1060–1068 (2010).

[57] Moore, R. J. et al. Fictrac: A visual method for tracking spherical motion and generating fictive animal paths. Journal of Neuroscience Methods 225, 106–119 (2014).

[58] Azevedo, A. W. et al. A size principle for recruitment of Drosophila leg motor neurons. eLife 9 (2020).

[59] Fuchs, E., Holmes, P., Kiemel, T. & Ayali, A. Intersegmental coordination of cockroach locomotion: adaptive control of centrally coupled pattern generator circuits. Frontiers in neural circuits 4, 125 (2011).

[60] Mantziaris, C. et al. Intra-and intersegmental influences among central pattern generating networks in the walking system of the stick insect. Journal of neurophysiology 118, 2296–2310 (2017).

[61] Schilling, M. & Cruse, H. Decentralized control of insect walking: A simple neural network explains a wide range of behavioral and neurophysiological results. PLoS computational biology 16 (2020).

[62] Ijspeert, A. J., Crespi, A., Ryczko, D. & Cabelguen, J.-M. From Swimming to Walking with a Salamander Robot Driven by a Spinal Cord Model. Science 315, 1416–1420 (2007).

[63] Ekeberg, Ö. A combined neuronal and mechanical model of fish swimming. Biological cybernetics 69, 363–374 (1993).

[64] Daun-Gruhn, S. A mathematical modeling study of inter-segmental coordination during stick insect walking. Journal of computational neuroscience 30, 255–278 (2011).

[65] DeAngelis, B. D., Zavatone-Veth, J. A. & Clark, D. A. The manifold structure of limb coordination in walking Drosophila. Elife 8, e46409 (2019).

[66] Oliveira, M., Santos, C. P., Costa, L., Matos, V. & Ferreira, M. Multi-objective parameter CPG optimization for gait generation of a quadruped robot considering behavioral diversity. In 2011 IEEE/RSJ International Conference on Intelligent Robots and Systems, 2286–2291 (2011).

[67] Deb, K., Pratap, A., Agarwal, S. & Meyarivan, T. A fast and elitist multiobjective genetic algorithm: NSGA-II. IEEE transactions on evolutionary computation 6, 182–197 (2002).

[68] Strauss, R. & Heisenberg, M. Coordination of legs during straight walking and turning in drosophila melanogaster. Journal of Comparative Physiology A 167 (1990).

[69] Takahashi, H. et al. Maximum force capacity of legs of a fruit fly during landing motion. In 19th International Conference on Solid-State Sensors, Actuators and Microsystems, 1061–1064 (2017).

[70] Elliott, C. J. & Sparrow, J. C. In vivo measurement of muscle output in intact Drosophila. Methods 56, 78–86 (2012).

[71] Mamiya, A., Gurung, P. & Tuthill, J. C. Neural coding of leg proprioception in Drosophila. Neuron 100, 636–650 (2018).

[72] Vincent, J. F. & Wegst, U. G. Design and mechanical properties of insect cuticle. Arthropod structure & development 33, 187–199 (2004).

[73] Flynn, P. C. & Kaufman, W. R. Mechanical properties of the cuticle of the tick Amblyomma hebraeum (Acari: Ixodidae). Journal of Experimental Biology 218, 2806–2814 (2015).

[74] Kimura, K.-i., Minami, R., Yamahama, Y., Hariyama, T. & Hosoda, N. Framework with cytoskeletal actin filaments forming insect footpad hairs inspires biomimetic adhesive device design. Communications biology 3, 1–7 (2020).

[75] Kuan, A. T. et al. Dense neuronal reconstruction through X-ray holographic nano-tomography. Nature neuroscience (2020).

[76] Chaffey, N. Principles and techniques of electron microscopy: biological applications (2001).

[77] Schindelin, J. et al. Fiji: an open-source platform for biological-image analysis. Nature methods 9, 676–682 (2012).

[78] van der Walt, S. et al. scikit-image: image processing in Python. PeerJ 2 (2014).

[79] Foundation, B. Blender-a 3D Modelling and Rendering Package (2012).

[80] Ferris, G. External morphology of the adult. Biology of Drosophila 368–419 (1950).

[81] Dickson, W. B., Straw, A. D. & Dickinson, M. H. Integrative Model of Drosophila Flight. AIAA Journal 46, 2150–2164 (2008).

[82] Geurten, B. R. H., Jôhde, P., Corthals, K. & Göpfert, M. C. Saccadic body turns in walking Drosophila. Frontiers in Behavioral Neuroscience 8, 365 (2014).

[83] Szczecinski, N. S., Bockemühl, T., Chockley, A. S. & Büschges, A. Static stability predicts the continuum of interleg coordination patterns in Drosophila. Journal of Experimental Biology 221 (2018).

[84] Mantziaris, C., Bockemühl, T. & Büschges, A. Central pattern generating networks in insect locomotion. Developmental neurobiology 80, 16–30 (2020).

[85] Cohen, A. H., Holmes, P. J. & Rand, R. H. The nature of the coupling between segmental oscillators of the lamprey spinal generator for locomotion: A mathematical model. Journal of Mathematical Biology 13, 345–369 (1982).

[86] Benitez-Hidalgo, A., Nebro, A. J., Garcia-Nieto, J., Oregi, I. & Del Ser, J. jMetalPy: A Python framework for multi-objective optimization with metaheuristics. Swarm and Evolutionary Computation 51 (2019).

